# A single-cell survey of *Drosophila* blood

**DOI:** 10.1101/2019.12.20.884999

**Authors:** Sudhir Gopal Tattikota, Yanhui Hu, Yifang Liu, Bumsik Cho, Victor Barrera, Michael Steinbaugh, Sang-Ho Yoon, Aram Comjean, Fangge Li, Franz Dervis, Ruei-Jiun Hung, Jin-Wu Nam, Shannan Ho Sui, Jiwon Shim, Norbert Perrimon

**Affiliations:** Department of Genetics, Blavatnik Institute, Harvard Medical School, Boston, USA; Department of Life Science, Hanyang University, Seoul, Korea; Harvard TH Chan Bioinformatics, Boston, USA; Howard Hughes Medical Institute, Boston, USA

**Keywords:** Drosophila, blood, hemocytes, cell states, scRNA-seq, wounding, wasp infestation, immune response

## Abstract

*Drosophila* blood cells, called hemocytes, are classified into plasmatocytes, crystal cells, and lamellocytes based on the expression of a few marker genes and cell morphologies, which are inadequate to classify the complete hemocyte repertoire. Here, we used single-cell RNA sequencing (scRNA-seq) to map hemocytes across different inflammatory conditions in larvae. We resolved plasmatocytes into different states based on the expression of genes involved in cell cycle, antimicrobial response, and metabolism together with the identification of intermediate states. Further, we discovered rare subsets within crystal cells and lamellocytes that express fibroblast growth factor (FGF) ligand *branchless* and receptor *breathless*, respectively. We demonstrate that these FGF components are required for mediating effective immune responses against parasitoid wasp eggs, highlighting a novel role for FGF signaling in inter-hemocyte crosstalk. Our scRNA-seq analysis reveals the diversity of hemocytes and provides a rich resource of gene expression profiles for a systems-level understanding of their functions.

**Highlights:**

- scRNA-seq of *Drosophila* blood recovers plasmatocytes, crystal cells, and lamellocytes
- scRNA-seq identifies different plasmatocyte states based on the expression of genes involved in cell cycle regulation, antimicrobial response, and metabolism
- Pseudotemporal ordering of single cells identifies crystal cell and lamellocyte intermediate states
- scRNA-seq uncovers a novel role for FGF signaling in inter-hemocyte crosstalk

## Introduction

The immune system forms an important layer of defense against pathogens in a wide variety of organisms including *Drosophila* (Banerjee et al., 2019; Mathey-Prevot and Perrimon, 1998). The chief mode of immune response in flies involves innate immunity, which is composed of diverse tissue types including fat body, gut, and blood cells called the hemocytes (Buchon et al., 2014). Hemocytes represent the myeloid-like immune cells, but so far have been considered less diverse compared to their vertebrate counterparts (Evans et al., 2003; Wood and Martin, 2017). In addition to progenitor cells or prohemocytes, three major types of hemocytes are known in *Drosophila*: plasmatocytes (PM), crystal cells (CC), and lamellocytes (LM). PMs are macrophage-like cells with parallels to vertebrate tissue macrophages, while CCs and LMs perform functions analogous to clotting and granuloma formation in vertebrates (Buchon et al., 2014). Hemocytes in the larva derive from two lineages: the lymph gland and the embryonic lineage, which in the larva forms resident (sessile) clusters of hemocytes in subepidermal locations, also known as hematopoietic pockets (Gold and Brückner, 2015; Holz et al., 2003; Jung, 2005; Makhijani et al., 2011). Prohemocytes can give rise to all mature hemocytes in the lymph gland (Banerjee et al., 2019). Likewise, the embryonic lineage, which consists of self-renewing PMs, is capable of producing CCs and LMs during inflammatory conditions (Gold and Brückner, 2015; Leitão and Sucena, 2015; Márkus et al., 2009). Whereas PMs are important for phagocytosis and represent ∼90-95% of total hemocytes, CCs, which constitute ∼5%, are required for wound healing and melanization (Banerjee et al., 2019; Evans et al., 2003). Importantly, CCs and LMs are required for survival upon injury or immunogenic conditions in *Drosophila*, highlighting the significance of these specialized blood cell types (Galko and Krasnow, 2004; Rizki, 1962).

Traditionally, the classification of hemocytes is based on two major criteria: cell morphology (Rizki, 1957, 1962; Shrestha and Gateff, 1982) and expression of a few marker genes (Evans et al., 2003, 2014; Kurucz et al., 2007). The paucity of markers available to define cell types and the low-resolution of cell morphologies may have hindered the identification of rare cell types and failed to distinguish transient states. For instance, a PM-like cell called the podocyte, which possibly corresponds to an intermediate state between PMs and LMs, has been reported but its transcriptional signature remains unknown (Honti et al., 2010; Rizki, 1957; Stofanko et al., 2010). Moreover, ultrastructural and microscopic evidence has also suggested that several subsets within PMs and CCs exist, but they have not been characterized at the molecular level (Rizki, 1957; Shrestha and Gateff, 1982). Finally, little is known about hemocyte lineage trajectories with regards to the source cells or precursors, or about intermediate states that exist on the path to terminal differentiation of mature cell types. Hence, it is important to thoroughly characterize the molecular signatures of all the dynamic states of mature cell types in steady state and inflammatory conditions.

Advances in single-cell RNA sequencing (scRNA-seq) technologies allow comprehensive characterization of complex tissues, including blood (Satija and Shalek, 2014). In particular, scRNA-seq is powerful not only for identifying cell types but also resolving cell states and their dynamic gene expression patterns that are often buried in bulk RNA measurements (Trapnell, 2015). For example, recent studies using various scRNA-seq platforms have helped identify novel subtypes within monocytes and dendritic cells (Villani et al., 2017) and activated states of T cells (Szabo et al., 2019) in human blood. Further, scRNA-seq has documented the continuous spectrum of differentiation along the hematopoietic lineage in various species (Macaulay et al., 2016; Nestorowa et al., 2016; Velten et al., 2017; Zhang et al., 2018). In addition, scRNA-seq data allows pseudotemporal ordering of cells to re-draw developmental trajectories of cellular lineages (Cao et al., 2019). Thus, scRNA-seq, in conjunction with various lineage trajectory algorithms, allows the precise characterization of 1. differentiated cell types and their subtypes, 2. transient intermediate states, 3. progenitor or precursor states, and 4. activated states, which are often influenced by mitotic, metabolic, or immune-activated gene modules (Adlung and Amit, 2018; Trapnell, 2015; Wagner et al., 2016).

Here, we performed scRNA-seq of *Drosophila* hemocytes in unwounded, wounded, and parasitic wasp infested larvae to comprehensively distinguish mature cell types from their transient intermediate states. Our scRNA-seq analysis identifies novel marker genes to existing cell types and distinguishes activated states within PMs enriched in various genes involved in the regulation of cell cycle, metabolism, and antimicrobial response. In addition, we could precisely distinguish mature CCs and LMs from their respective intermediate states. Interestingly, our scRNA-seq revealed the expression of fibroblast growth factor (FGF) receptor *breathless* (*btl*) and its ligand *branchless* (*bnl*), in rare subsets of LMs and CCs, respectively, which we implicate in regulating effective immune responses against parasitoid wasp eggs *in vivo*. Altogether, our scRNA-seq analysis documents the diversity of hemocyte cell populations circulating in the fly blood and provides a resource of gene expression profiles of the various cell types and their states in *Drosophila*. This resource can be mined using a user-friendly searchable web-tool (www.flyrnai.org/scRNA/blood/) where genes can be queried, visualized, and compared across conditions.

## Results

### scRNA-seq of *Drosophila* hemocytes

Hemocyte differentiation can be induced in *Drosophila* larvae by mechanical wounding or oviposition by wasps such as *Leptopilina boulardi* (Rizki and Rizki, 1992). Hence, to characterize hemocyte populations and their heterogeneity, we first performed the two immune responsive conditions: wounded and wasp 24h post-infested (wasp inf. 24h), together with unwounded (Unw) control conditions (Fig. 1A). Further, to mobilize the sessile hemocytes into circulation, we briefly vortexed the larvae prior to bleeding (Petraki et al., 2015).

**Figure 1:**
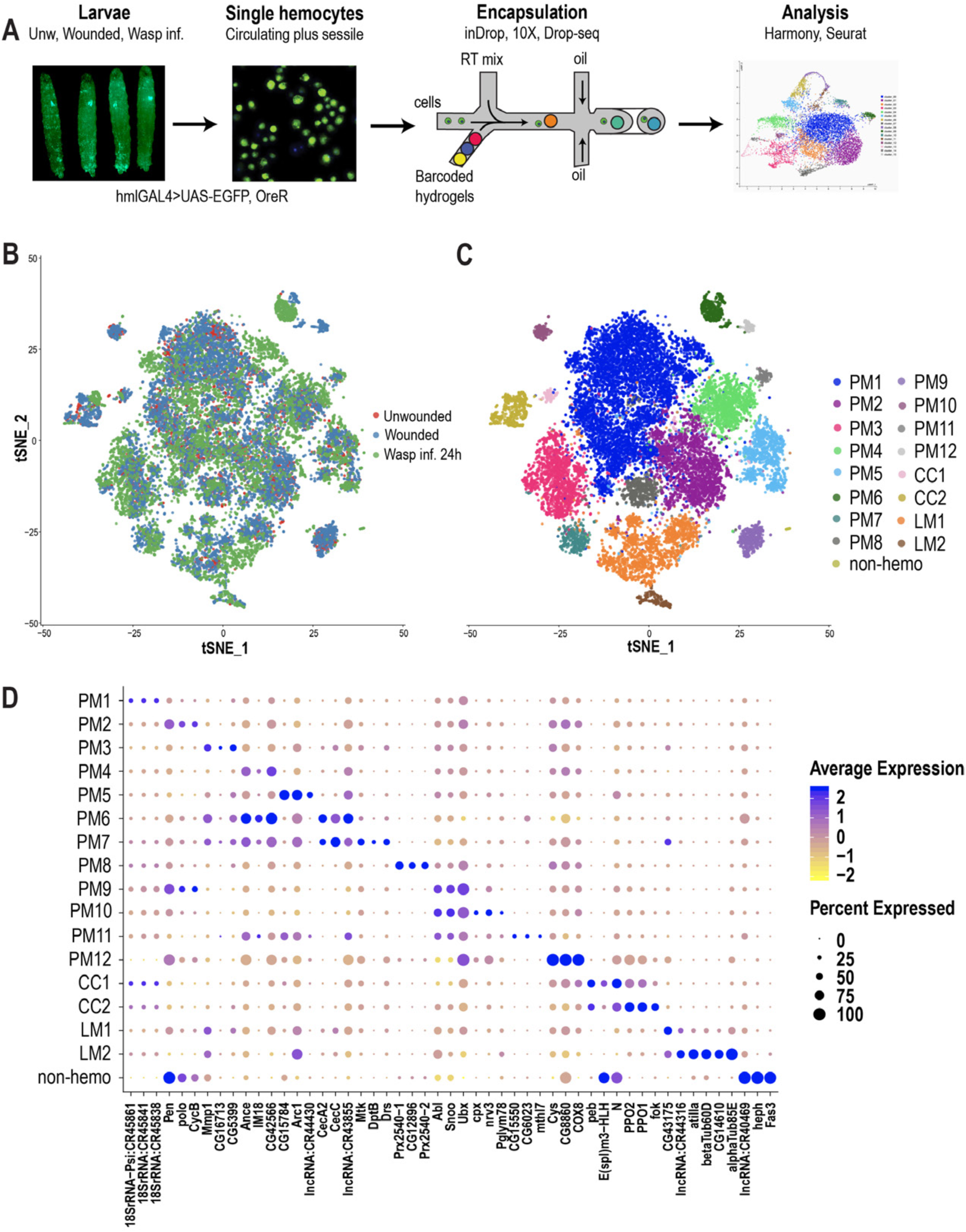
scRNA-seq of *Drosophila* hemocytes. A) Schematic of the scRNA-seq workflow. B) t-Distributed Stochastic Neighbor Embedding (t-SNE) plot of Harmony-based batch correction and integration of Unw (red), Wounded (blue), and Wasp inf. 24h (green) data sets. C) Clustering of batch corrected cells from all three conditions reveals a total of 17 clusters. D) Dot plot representing the top three genes enriched per cluster based on average expression (avg_logFC). Color gradient of the dot represents the expression level, while the size represents percentage of cells expressing any gene per cluster.

Subsequently, single hemocytes were encapsulated using microfluidics-based scRNA-seq technologies including inDrops (Klein et al., 2015), 10X Chromium (Zheng et al., 2017) or Drop-seq (Macosko et al., 2015). A total of 19,458 cells were profiled, with 3-4 replicates per condition, and obtained a median of 1010 genes and 2883 unique molecular identifiers (UMIs) per cell across all conditions (Table S1; Fig. S1A, B). In order to achieve a comprehensive map of all the hemocytes profiled by the three scRNA-seq platforms, we merged all data sets. We observed notable “batch effects” where cell types were being clustered according to condition, replicate, or technology (Fig. S1C, D, E). Thus, we applied the Harmony batch correcting method (Korsunsky et al., 2019), which is integrated into the Seurat R package (Stuart et al., 2019). Harmony successfully integrated all three data sets, including their replicates (Fig. 1B; S1D’, E’), and identified a total of 17 clusters (Fig. 1C). Based on known marker genes, we confidently assigned certain clusters as plasmatocytes (PMs marked by *Hemolectin* [*Hml*]), crystal cells (CCs marked by *lozenge* [*lz*]), or lamellocytes (LMs marked by *Atilla*) (Fig. 1C; S1G). Of the 17 clusters, one small cluster, representing ∼0.2% of the total profiled cells, did not express any of the pan-hemocyte markers such as *Hemese* (*He*) and *Serpent* (*srp*) (Fig. S1G) (Evans et al., 2014). Hence, we labeled this cluster as non-hemocyte (non-hemo).

### Diversity of hemocyte populations and their transcriptional dynamics

#### PM clusters

Despite the fact that PMs constitute over 90-95% of the total hemocyte pool, subclasses within this major cell type have not been described. The majority of the clusters we identified (12/17) express *Hml* but not CC or LM markers and thus we annotated them as PM clusters: PM1-12 (Fig. 1C; S1G).

PM1 represents the largest cluster and is enriched in rRNA genes such as *18SrRNA:CR45841* (Fig. 1D). Interestingly, there is evidence in mammals that certain rRNA genes, including pre-rRNA molecules, accumulate in activated macrophages (Radzioch et al., 1987; Varesio, 1985). We determined PM2 as the cycling or self-renewing state of PMs based on the expression of genes related to cell cycle such as *CycB*, *stg*, and *polo*, which are markers of G2/M stages of the cell cycle (Edgar and O’Farrell, 1990; Glover, 2005; Whitfield et al., 1990). Next, we identified a group of PM clusters, PM3-5, which are enriched in several immune-induced genes, including *Matrix metalloproteinase 1* (*Mmp1*) and *Immune induced molecule 18* (*IM18*). Further, our scRNA-seq identified two distinct clusters, PM6 and 7, expressing several genes that encode antimicrobial peptides (AMP) (Fig. 1D). PM6 highly expresses genes of the cecropin family, including *CecA2* and *CecC*, while PM7 is enriched in additional AMPs such as *Mtk*, *DptB*, and *Drs* (Fig. 1D). The differential expression of this broad spectrum of AMPs (Hoffmann and Reichhart, 2002; Tzou et al., 2002) in PM6 and 7 (collectively termed PM^AMP^) suggests that hemocytes can elicit a humoral immune response against a variety of pathogens. Together, we define PM3-7 as the immune-activated states of the PMs because of the expression of several immune-induced genes including the AMP-genes.

In addition to these major clusters, our scRNA-seq also identified several minor clusters: PM8-12. PM8 is enriched in genes encoding peroxidase enzymes such as *Prx2540-1*, *Prx2540-2*, and *CG12896*, which is an uncharacterized gene highly similar to *peroxiredoxin 6* (*PRDX6*) in humans. Although the function of these genes is not known in the context of hemocytes and immunity, peroxisomes have been reported to be necessary for phagocytosis by macrophages in both mice and *Drosophila* (Di Cara et al., 2017). PM9 and PM10 are enriched in the protooncogenes *Abl*, *Sno oncogene* (*Snoo*), and the transcriptional regulator *Ultrabithorax* (*Ubx*). PM11 and PM12 represent the smallest clusters within PMs and are defined by the expression of the uncharacterized genes *CG15550* and *CG6023*, together with a methuselah-type receptor gene, *methuselah-like 7* (*mthl7*) in PM11, and *Cys*, *CG8860*, and *COX8* in PM12 (Fig. 1D; Table S2).

#### CC clusters

The next most abundant immune responsive cells are the CCs, which are the main source of two enzymes important for melanization, prophenoloxidase 1 and 2 (PPO1 and PPO2). These enzymes are critical for survival upon wounding in larvae and adults (Binggeli et al., 2014; Dudzic et al., 2019; Theopold et al., 2014). Based on the expression of *PPO1* and *PPO2* together with the gene encoding the Runt related transcription factor *Lz*, two clusters were assigned to CCs: CC1 and CC2 (Fig. 1C-D; S1G). CC1 expresses low levels of *PPO1* but high levels of *Notch*, *pebbled* (*peb*), and the enhancer of split complex gene *E(spl)m3-HLH* (Fig. 1D). Although *Notch* and *peb* have been shown to be associated with CC development (Terriente-Felix et al., 2013), expression of the Notch target gene *E(spl)m3-HLH* (Couturier et al., 2019) has not been reported. On the other hand, CC2 shows higher expression levels of *PPO1* and *PPO2* genes. Hence, we consider CC2 to represent mature CCs, while CC1 may represent an immature or a transient intermediate state.

#### LM clusters

LMs represent the rarest cell type, the numbers of which dramatically increase during wounding and wasp infestation (Márkus et al., 2005; Rizki and Rizki, 1992). Based on the expression of the LM marker gene *Atilla* (Evans et al., 2014; Kurucz et al., 2007), we assigned two clusters to LMs: LM1 and LM2 (Fig. 1C-D; S1G). LM1 is the larger cluster of the two and is enriched in *Atilla* besides a long non-coding RNA, *lncRNA:CR44316* (Fig. 1D). LM2 represents a smaller cluster expressing *Atilla*, *betaTub60D*, and *alphaTub85E*. The expression level of *Atilla* is higher in LM2 compared to LM1 (Fig. 1D), suggesting that LM2 represents mature LMs, whereas LM1 may represent the LM intermediate state. Moreover, the strong expression pattern of tubulins and other cytoskeletal proteins may be important for the maintenance of structural integrity of LMs and their dynamic roles in encapsulation (Rizki and Rizki, 1994).

Altogether, our scRNA-seq analysis recovered all major cell types within the hemocyte repertoire including the fine-grained dissection of PMs into self-renewing or cell-cycle (PM2) and various immune-activated states (PM3-7). The functions of the newly identified genes in the rest of the PM clusters, which are minor subpopulations except for PM1, remain to be characterized. Presumably, these subpopulations represent transiting intermediates along the course of terminal differentiation or activated states of PMs and other cell types. We also identified two clusters each for CCs and LMs, which display differential expression of their marker genes, *PPO1* and *Atilla*, respectively (Fig. 1D; see Table S2 for additional marker genes).

### Pseudotemporal ordering of cells delineates hemocyte lineages

PMs of the embryonic lineage reside in larval hematopoietic pockets and, over the course of third larval instar, increasingly enter the hemolymph to circulate in the open circulatory system (Makhijani et al., 2011).

The generation of these PMs, initially by differentiation from embryonic progenitors, and later in the larva through self-renewal of differentiated PMs, is well established (Makhijani et al., 2011). In contrast, the development of terminally differentiated CCs and LMs from the embryonic lineage has remained speculative, but includes models of transdifferentiation from PMs (Anderl et al., 2016; Leitão and Sucena, 2015; Márkus et al., 2009). Since hemocyte progenitors reside only in the lymph gland and are absent in circulation or sessile compartments (Jung et al., 2005), it is important to address the immediate sources of mature cell types in circulation. scRNA-seq data can be used to construct pseudotemporal relationships between individual cell transcriptomes and impute cell lineages derived from precursor cells (Trapnell, 2015; Trapnell et al., 2014). We took advantage of our observation that PM2 expresses cell cycle genes to construct lineage trees emerging from this cell-cycle state. To avoid using batch corrected cells, we chose 10X genomics-derived wounded data set (Fig. S2A; Table S1), which represents all the mature cell types. We used Monocle3 (Cao et al., 2019) and assigned PM2 as the start point of the pseudotime intervals (Fig. 2A-B; S2A-C). Monocle3 data shows that three major lineages emerge from the start point (Fig. 2C-E). Lineage1 terminates in CC fate, and includes the two CC clusters, CC1 and CC2. As expected, CC1 precedes CC2, strongly supporting that CC1 represents an intermediate state (CC^int^) (Fig. 2B). Consistently, the expression of *PPO1* steadily increases from CC1 and reaches its peak level upon becoming mature CCs (Fig. 2F). Lineage2 on the other hand terminates in a fate that include PM1 and other minor PM clusters (PM8-12) (Fig. 2B, D). Lastly, Lineage3 leads towards the clusters of the immune-activated state together with mature LMs (Fig. 2B, E). As expected, LM1 precedes LM2, supporting our initial observation that LM1 corresponds to the intermediate state of LMs (LM^int^). This is further demonstrated by the expression of *Atilla*, which steadily increases with low levels in LM1 and higher levels in LM2 (Fig. 2G). Lineage3 also terminates with cells of the immune-activated state, PM7 and PM5 (Fig. 2B), which express *Drs* (in PM7) and genes pertaining to glycolysis and GSTs (in PM5) (Fig. 2H; S2F-H).

**Figure 2:**
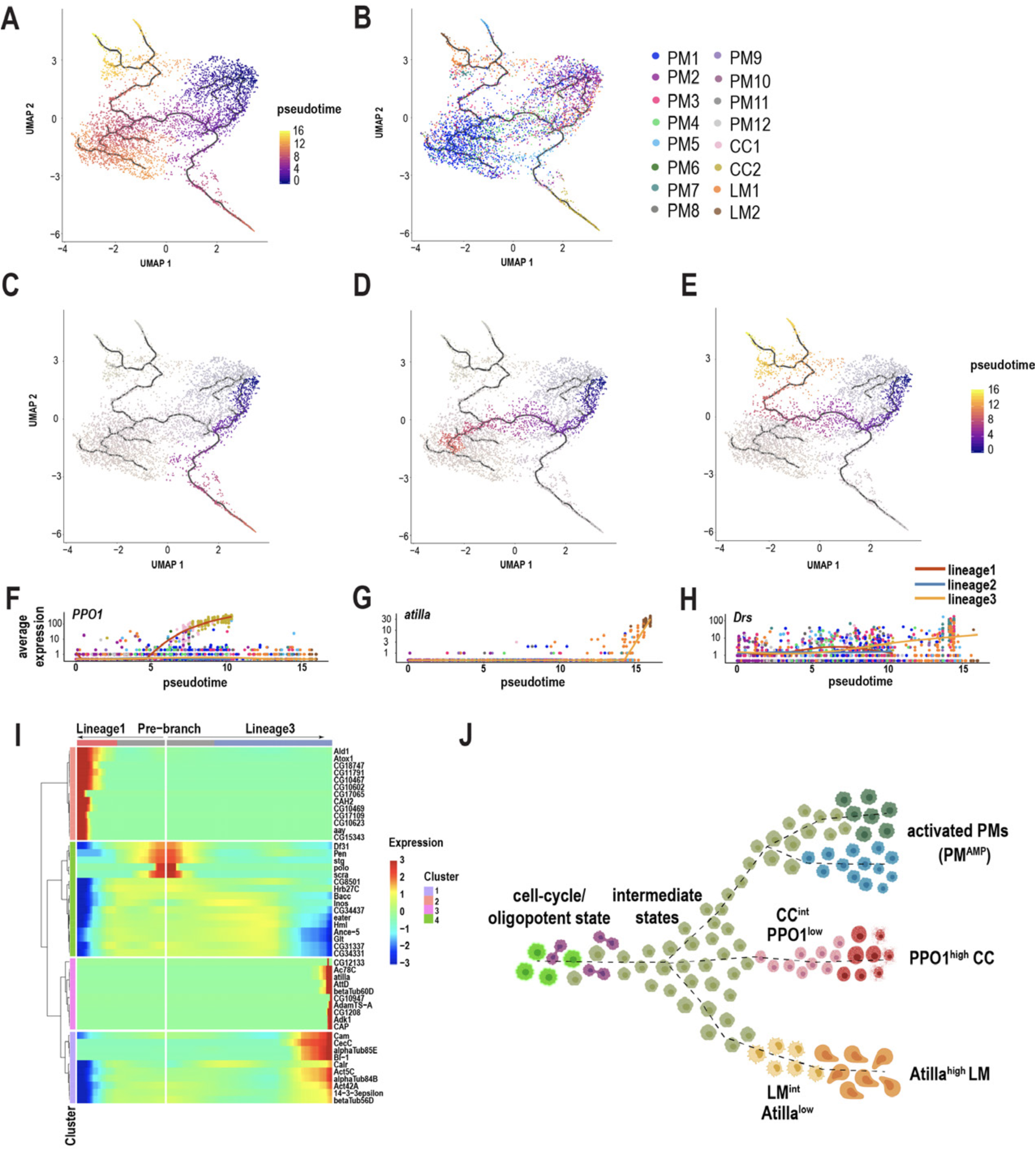
Pseudotemporal ordering of cells delineates immune cell lineages. A) Monocle3 was used to track cells over pseudotime on the 10X-derived wounded data set. B) Visualization of clusters (from Fig. 1C) onto the pseudotime map. C-E) Three major lineage routes were obtained from the start site. F-H) Mapping gene expression over pseudotime. Expression of *PPO1* (F), *Atilla* (G), and *Drs* (H). I) Monocle-based gene expression signature between the Lineages 1 and 3 with the ‘pre-branch’ in the middle. J) Schematic showing potential lineage flow from PM2 to mature cell types with their intermediates.

Next, to gain deeper insights into the gene expression signatures along the pseudotime intervals, we analyzed the differentially expressed genes (DEG) between the source and Lineage1 or Lineage3. DEG analysis over pseudotime revealed four major clusters depending on the expression of marker genes at the beginning (pre-branch) and end of the pseudotime interval along Lineage1 and 3 (Fig. 2I). Interestingly, the pre-branch is enriched with PM marker genes such as *eater* and *Hml*, suggesting that precursor cells are indeed the self-renewing PMs. Moreover, the expression of the cell cycle genes gradually decrease as the lineages progress towards 1 or 3 (Fig. 2I), consistent with a relationship between cell cycle arrest and terminal differentiation in vertebrates (Morse et al., 1997; Ruijtenberg and van den Heuvel, 2016; Soufi and Dalton, 2016). To assess whether blocking the cell cycle promotes terminal differentiation, we expressed RNAi against one of the top enriched cell cycle genes, *polo*, in Hml^+^ PMs (*HmlGAL4>polo-i*). RNAi-mediated knockdown of *polo* resulted in a significant increase in the production of LMs compared to controls (Fig. S2I-K), suggesting that cell cycle arrest may be required for terminal differentiation of cell types. Finally, to visualize cells accumulating along the pseudotime intervals, we analyzed the cell densities using ridge plots, which revealed that PM8-12 accumulated predominantly between the start and end of pseudotime intervals, indicating that they are all transient PM intermediates (PM^int^) (Fig. S2D).

Altogether, based on Monocle3, we propose that PM2 has oligopotent potential and can give rise to terminally differentiated cell types and possibly other activated states within PMs. Further, our analysis confirms the existence of CC and LM intermediate states that precede their fully differentiated mature cell types (Fig. 2J).

### Changes in hemocyte composition and identification of a novel Mtk-like AMP

In addition to identifying genes enriched in each cluster and their changes across conditions, it is possible to estimate cell fraction changes from scRNA-seq data sets. To achieve this, we first segregated the three conditions (Fig. 3A-C), then calculated cell fraction changes. Whereas PM1-5, 8, and 11 were well represented in all three conditions, PM9 is negligibly detected in the wasp inf. 24h condition and PM10 and 12 were majorly detected in the wounded condition. The PM^AMP^ clusters, PM6 and 7, emerged mainly upon wounding or wasp infestation compared to Unw controls (Fig. 3A-D). Strikingly, both CC1 and CC2 were underrepresented in wasp inf. 24h, consistent with previous observations that CC numbers dramatically decrease following oviposition by wasps (Kacsoh and Schlenke, 2012) (Fig. 3A-D). Of the LM clusters, LM1 was detected in all three conditions, however, LM2 emerged only upon wounding or wasp infestation. In summary, LM2 and PM^AMP^ are the major clusters that are represented mostly upon wounding or wasp infestation.

**Figure 3:**
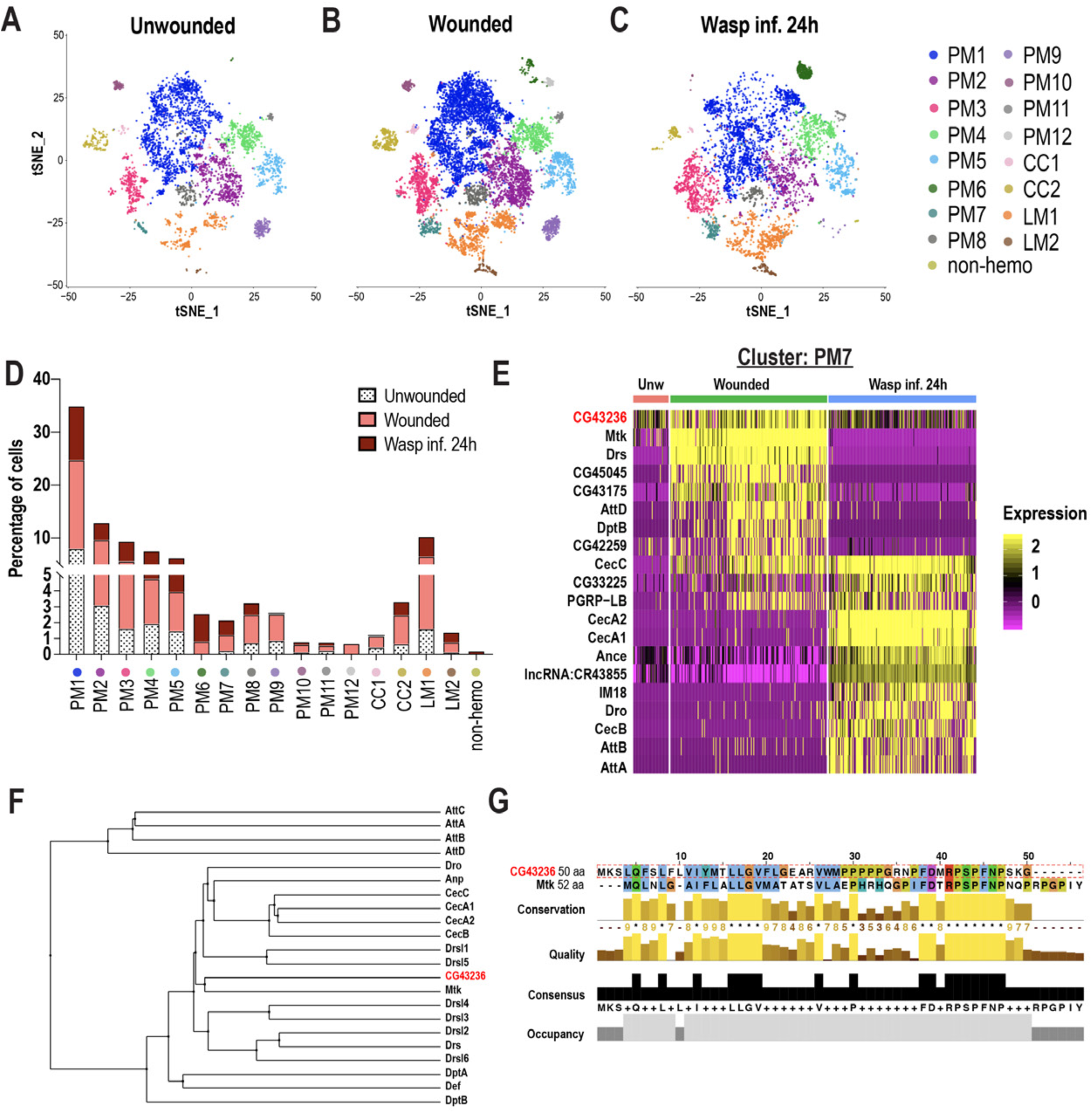
Changes in immune cell composition and identification of a novel Mtk-like AMP. A-C) t-SNE plots of A) Unwounded (Unw), B) Wounded, and C) Wasp inf. 24h conditions. D) Cell fraction changes in clusters based on treatment conditions. E) Heat map profile of the top expressed genes in cluster PM7. Genes were ranked based on expression levels in each condition in the heat map. F) Phylogenetic tree map constructed with the peptide sequences of all known AMPs together with CG43236. G) Global alignment of CG43236 and Mtk peptide sequences using Jalview protein alignment software.

Despite some PM clusters being represented in all three conditions, their gene expression patterns are specific to wounding or wasp inf. 24h. For instance, *Mmp1* showed increased expression only upon wounding or wasp inf. 24h in PM3 compared to Unw controls (Fig. S3A). Likewise, the increased expression of GST genes such as *GstE6* is specific to wounded or wasp inf. 24h (Fig. S3B). With regards to the PM6 cluster of immune-activated state, *CecA2* showed an increased expression specifically in wasp inf. 24h compared to wounded conditions (Fig. S3C). Further, DEG analysis of PM7 revealed that most of the AMP genes were expressed only during wounding or wasp inf. 24h and not in Unw controls (Fig. 3E). Of note, the AMP gene signature was unique to either wounded or wasp inf. 24h conditions. For example, whereas *Mtk*, *Drs*, *DptB*, and *AttD* were specific to wounded condition, *CecA1*, *CecB*, *Dro*, and *AttA* were unique to wasp inf. 24h (Fig. 3E). These data indicate that clusters represented in all three conditions may nevertheless differ from each other in a condition-specific manner with respect to their differential gene signatures (Table S3).

A survey of all top enriched genes in PM7 revealed the identification of an uncharacterized gene, *CG43236* (Fig. 3E). *CG43236* encodes a small peptide of 50 amino acids (aa) and its phylogenetic alignment with all known AMPs revealed that it clustered with Mtk, an antibacterial and antifungal AMP (Levashina et al., 1995, 1998) (Fig. 3F, G). Both Mtk and CG43236 possess an Antimicrobial10 domain that is unique to the Metchnikowin family (Fig. 3G; S3D-D’), leading us to name *CG43236* as *Mtk-like (Mtkl*). To validate its expression within hemocytes and whole larvae, we performed qRT-PCR in Unw control and wounded conditions. Consistent with our scRNA-seq data (Fig. 3E), *CG43236* is well expressed in Unw control hemocytes based on average Ct values compared to *Rp49* (Fig. S3E). However, the induction of *CG43236* was modest, if any, compared to the robust induction of *CecC*, *Drs*, and *Mtk* upon wounding in hemocytes (Fig. S3F), suggesting that *CG43236* may be strongly regulated by additional modes of wounding such as sepsis or fungal infection. However, its expression was strongly induced upon wounding in whole larvae, indicating that *CG43236* may also be expressed by non-hemocytic cells (Fig. S3G). Finally, the expression levels of both *Mtk* and *CG43236* gradually increase in Lineage3 over pseudotime intervals, indicating their specificity to this lineage where PM7 terminates (Fig. 3H, I). Altogether, the DEG analysis of PM7 identified a novel Mtk-like AMP.

### Crystal cell sub-clustering distinguishes CC intermediates from mature CCs

Because CCs split into two distinct clusters (CC1 and CC2) (Fig. 1C), we sub-clustered these cells independently of the other clusters. We used Harmony to correct for batch effects arising from the technological platforms and conditions (Fig. S4A, B). Subsequent cell clustering revealed two distinct clusters: one with low *PPO1* expression (PPO1^low^) and one with very high *PPO1* expression (PPO1^high^) (Fig. 4A, C).

**Figure 4:**
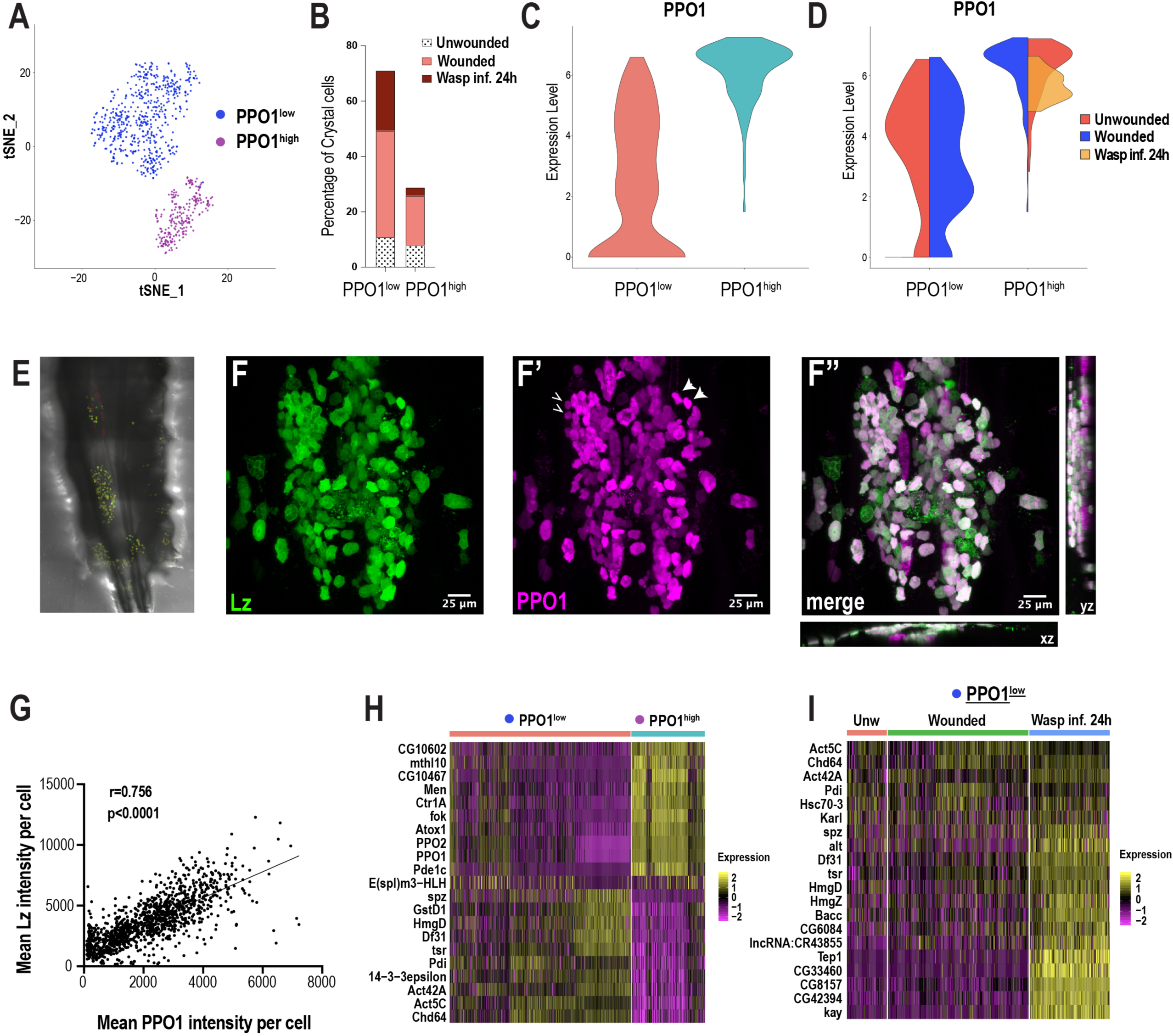
Crystal cell sub-clustering distinguishes CC intermediates from mature CCs. A) t-SNE plot of CC sub-clustering depicting two CC sub-clusters, PPO1^low^ and PPO1^high^. B) Percentage of PPO1^low^ and PPO1^high^ CCs across the three conditions. C) Violin plot indicating the expression level of *PPO1* in the two CC clusters. D) Expression level of *PPO1* across the three conditions. E) Confocal images of the posterior-dorsal side of a representative *LzGAL4; UAS-GFP, BcF6-mCherry* third instar larva. BcF6-mCherry is a PPO1 enhancer trap, which is a reporter for CCs. F-F’’) Confocal images of GFP+ (F), BcF6-mCherry+ (F’), and merged GFP+ mCherry+ CCs (F’’). xz and yz images in F’’ represent the depth of the stacks. Representative PPO1^low^ and PPO1^high^ CCs are shown by open and solid arrow heads, respectively, in F’. Scale bar = 25 μm. G) Mean intensities of GFP (Lz) and mCherry (PPO1). The correlation plot represents data from unwounded *LzGAL4; UAS-GFP, BcF6-mCherry* larvae (n=18; total CCs analyzed=1233). H-I) Heat maps of marker gene expression in PPO1^low^ and PPO1^high^ clusters (H) and differentially expressed gene (DEG) analysis of the marker genes across conditions in PPO1^low^ cluster (I).

While the percentage of PPO1^low^ CCs was higher than that of the mature PPO1^high^ CCs, the fraction of these cells increased upon wounding (Fig. 4B, S4C-E). Furthermore, the expression level of *PPO1* was similar between Unw controls and wounded condition in both clusters. Interestingly, its expression was negligibly detected in PPO1^low^ CCs and slightly lower in PPO1^high^ CCs in wasp infested larvae (Fig. 4D). To determine whether the two CC clusters coexist as distinct populations or whether PPO1^low^ is a CC^int^ state along the course of CC maturation, we used *LzGAL4, UAS-GFP; BcF6-mCherry* larvae to label Lz^+^ and PPO1^+^ CCs with GFP and mCherry, respectively. We examined the dorso-posterior end, where clusters of hemocytes that include CCs reside along the dorsal vessel of larvae (Fig. 4E) (Leitão and Sucena, 2015). In line with the scRNA-seq data, *in vivo* imaging analysis revealed distinct populations of CCs within the sessile hub, with CCs displaying differential intensities of GFP and mCherry (Fig. 4F-F’’). Furthermore, intensity measurements of GFP and mCherry revealed a significant positive correlation (Fig. 4G), which is consistent with previous studies demonstrating that Lz, together with Srp, can activate the expression of *PPO1* (Waltzer et al., 2003).

Of note, the correlation plot did not reveal two separate populations of CCs, but rather heterogenous cell populations, existing in a potential continuum (Fig. 4G). This further supports the Monocle3 prediction that PPO1^low^ may represent the CC^int^ state, whereas PPO1^high^ corresponds to mature CCs (Fig. 2B, F).

As noted above, PPO1^high^ CCs express many of the mature CC marker genes, including *PPO2* (Fig. 4H). This cluster also highly expresses a number of uncharacterized genes such as *CG10602* and *CG10467*, which potentially encode enzymes with epoxide hydrolase and aldose 1-epimerase activities, respectively (Fig. 4H). The human ortholog of *CG10602*, *LTA4H* (*leukotriene A4 hydrolase*), encodes an enzyme involved in the biosynthesis of a proinflammatory mediator, leukotriene B4 (Crooks and Stockley, 1998). Moreover, mutations in *lta4h* render zebrafish hypersusceptible to mycobacterial infections (Tobin et al., 2010). These observations suggest that *CG10602* may play an important role for mature CCs in combating bacterial infections, in addition to their role in melanization. On the contrary, PPO1^low^ CCs are enriched in *spatzle* (*spz*), a cytokine that activates the Toll pathway (Lemaitre et al., 1996) (Fig. 4H). Besides *spz*, PPO1^low^ CCs express cell cycle/chromatin associated genes such as the *Decondensation factor 31* (*Df31*) and *HmgD* (Fig. 4H), suggesting that PPO1^low^ may be in a cycling or proliferative state. Furthermore, although many of the genes, including *spz*, *Df31*, and *HmgD,* were more enriched upon wasp inf. 24h in PPO1^low^ (Fig. 4I), the PPO1^high^cluster did not display notable changes in gene expression (Fig. S4F). Of note, we confirmed the expression of a novel CC marker gene *E(spl)m3-HLH*, which is expressed in ∼13% of CCs in both PPO1^low^ and PPO1^high^ populations (Fig. S4G-G’’’). In addition, Monocle3 revealed an enrichment of *E(spl)m3-HLH* in the CC2 cluster in Lineage1 (the CC lineage) over the pseudotime intervals (Fig. S4H), suggesting that it is expressed early during CC development. In summary, CC sub-clustering identifies a CC intermediate state and highlights the possibility that CCs exist in a continuum, along the course of CC maturation, as suggested by Monocle3 and *in vivo* imaging data.

### Lamellocyte sub-clustering identifies LM intermediates and subtypes

Previous studies have speculated the presence of LM intermediates (Anderl et al., 2016; Honti et al., 2010; Rizki, 1957; Stofanko et al., 2010). Monocle3 predicted that LM1 might correspond to a LM intermediate state (Fig. 2B, G, J). To further test this prediction, we sub-clustered all the Atilla^+^ LMs derived from all conditions. To increase the diversity of LMs, we performed scRNA-seq at one additional time point of wasp infestation: wasp inf. 48h. Clustering analysis revealed that more than 50% of all cells are of the LM lineage (S5A-C). To sub-cluster the LMs, we considered the Atilla^+^ clusters 0 and 1 from wasp inf. 48h (Fig. S5A-B) together with Atilla^+^ LMs from Unw, wounded, and wasp inf. 24h data sets (Fig. 1C-D). We used Harmony to correct for batch effects (Fig. S5D-E) and subsequent clustering of all the LMs revealed 5 distinct clusters (Fig. 5A), that we named LM1-4 and CC based on the expression of top enriched genes (Fig. 5E). Cell fraction calculations revealed that all the clusters, especially LM4 and CC, were predominantly derived from wasp inf. 48h condition (Fig. 5B; S5E-I). The last cluster was annotated as CC based on extremely low or no expression of *Atilla* and enrichment of CC marker genes including *PPO1* (Fig. 5C, E). We speculate that LM1 may represent a LM^int^ state based on the low level of *Atilla* together with enrichment of PM marker genes such as *Pxn* and *Hml*, which are usually not expressed upon LM maturation (Fig. 2I; 5E) (Stofanko et al., 2010). Interestingly, expression of *Atilla* is negligible in Unw control and wounded conditions, while *Hml* is higher compared to wasp inf. conditions in LM1 (Fig. 5D; S5J).

**Figure 5:**
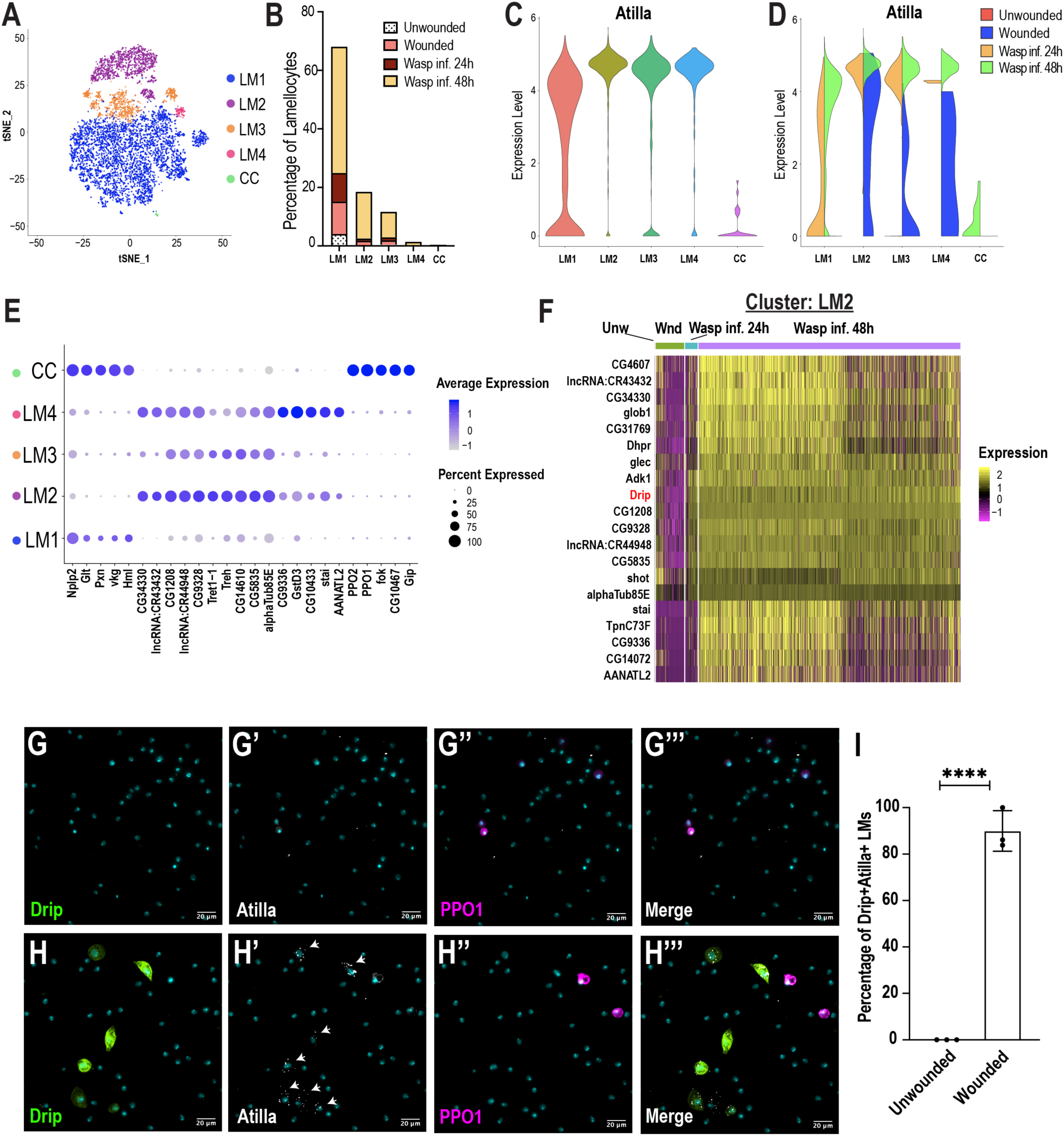
Lamellocyte sub-clustering identifies LM intermediates and subtypes. A) t-SNE plot depicting the LM sub-clusters. B) Changes in LM fractions across the three conditions. C) Expression of the LM marker gene *Atilla* was used to annotate the LM clusters. D) Split violin plot shows the differential expression of *Atilla* in different conditions. E) Dot plot representing the top 5 marker genes per LM cluster. F) Heat map depicts the DEG analysis of top genes in LM2 across all conditions. G-H’’’) Expression validation of *Drip* in hemocytes derived from *DripGAL4>mCD8-GFP; BcF6-mCherry* Unw (G-G’’’) or wounded (H-H’’’) larvae. Nuclei are stained with DAPI (Cyan). Scale bar = 20 μm. I) Percentage of GFP^+^ Atilla^+^ LMs normalized to total Atilla^+^ LMs per field of view. Data is represented by three independent biological replicates (n=3). The error bars are represented as ±SD (standard deviation). P values are represented by **** (p<0.0001).

Based on the high level of expression of *Atilla*, we conclude that LM2, LM3, and LM4 are subtypes of mature LMs (Fig. 5C). Whereas LM3 and LM4 expressed relatively low levels of *Trehalase* (*Treh*) and its transporter *Tret1-1*, LM2 displayed high levels of these genes (Fig. 5E). In addition to *Treh* and *Tret1-1*, almost all LM2 cells express two uncharacterized genes that potentially encode sugar transporters, *CG4607* and *CG1208* (Fig. 5E, F). These genes emerged as the top-expressed genes in wasp inf. 48h when compared to the Unw control, wounded, or wasp inf. 24h conditions (Fig. 5F). Of note, CG4607 and CG1208 belong to the SLC2 family of hexose sugar transporters and are orthologous to SLC2A6/GLUT6 or SLC2A8/GLUT8 in humans. Importantly, it has recently been demonstrated that GLUT6 acts as a lysosomal transporter, which is regulated by inflammatory stimuli (Maedera et al., 2019) and is strongly upregulated in macrophages activated by LPS (Caruana et al., 2019). Similarly, in *Drosophila*, immune activation is energy demanding and glucose may be used as a major source of energy for the production of mature LMs (Bajgar et al., 2015). These observations suggest that LMs may require specialized sugar transporters for their maturation to elicit an effective immune response against parasitoid wasp eggs.

In addition to the expression of the sugar transporters, all LM sub-clusters are enriched in a water transporter, Drip (AQP4 in humans), which has recently been shown to play a role in T-cell proliferation and activation in mice (Nicosia et al., 2019) (Fig. 5F; S5J-L). To confirm *Drip* expression *in vivo*, third instar *DripGAL4, UAS-mCD8-GFP; BcF6-mCherry* larvae were wounded and then monitored for the expression of *Drip*. As expected, Drip^+^ GFP cells were not detected during steady state in Unw control larvae (Fig. 5G-G’’’, 5I). However, Drip^+^ GFP cells, significantly increased upon wounding, and strikingly, the expression of Drip was detected in ∼90% of Atilla^+^ LMs (Fig. 5H-H’’’, 5I). Finally, the smaller clusters of LMs, LM3 and LM4, were also enriched in genes similar to those expressed in LM2, albeit at a lower level (Fig. 5E), suggesting that LM3-4 are subtypes of mature LMs. In contrast to LM3, LM4 expresses additional marker genes that include *GstD3* suggesting that this subtype of LMs may also be involved in the detoxification of xenobiotics (Fig. 5E; S5L). In summary, LM sub-clustering identified the LM^int^ state together with novel marker genes specific to different subsets of mature LMs.

### scRNA-seq uncovers a novel role for the FGF pathway in immune response

To identify signaling pathways enriched in each cluster, we performed pathway enrichment analysis on the scRNA-seq data across all conditions (Fig. 6A). As expected, the PM6 and PM7 clusters are enriched in Imd and to lesser extent Toll signaling pathways (Lemaitre and Hoffmann, 2007). CC1 and CC2 are highly enriched in Notch signaling components (Fig. 6A; Table S4), which is consistent with previous studies that showed the importance of Notch in CC development (Duvic et al., 2002; Leitão and Sucena, 2015). Further, PM3 was highly enriched in particular for components of the TNF-alpha, Imd, and Toll pathways. In addition to these known pathways, we also identified the fibroblast growth factor (FGF) signaling pathway to be highly enriched in the LMs (Fig. 6A; Table S4). Although the fly FGFR, Heartless (Htl) and its ligands, Thisbe (Ths) and Pyramus (Pyr) have been shown to be required for progenitor differentiation in the lymph gland (Dragojlovic-Munther and Martinez-Agosto, 2013), the role of FGF signaling has not been addressed in circulating hemocytes of the embryonic lineage. Interestingly, the second FGFR gene, *breathless* (*btl*), is one of the components that is enriched albeit at low levels in LM2 (Fig. 6B, B’). In addition, a small subpopulation of CCs expresses *branchless* (*bnl*), which encodes the only ligand for Btl (Fig. 6B, B’).

**Figure 6:**
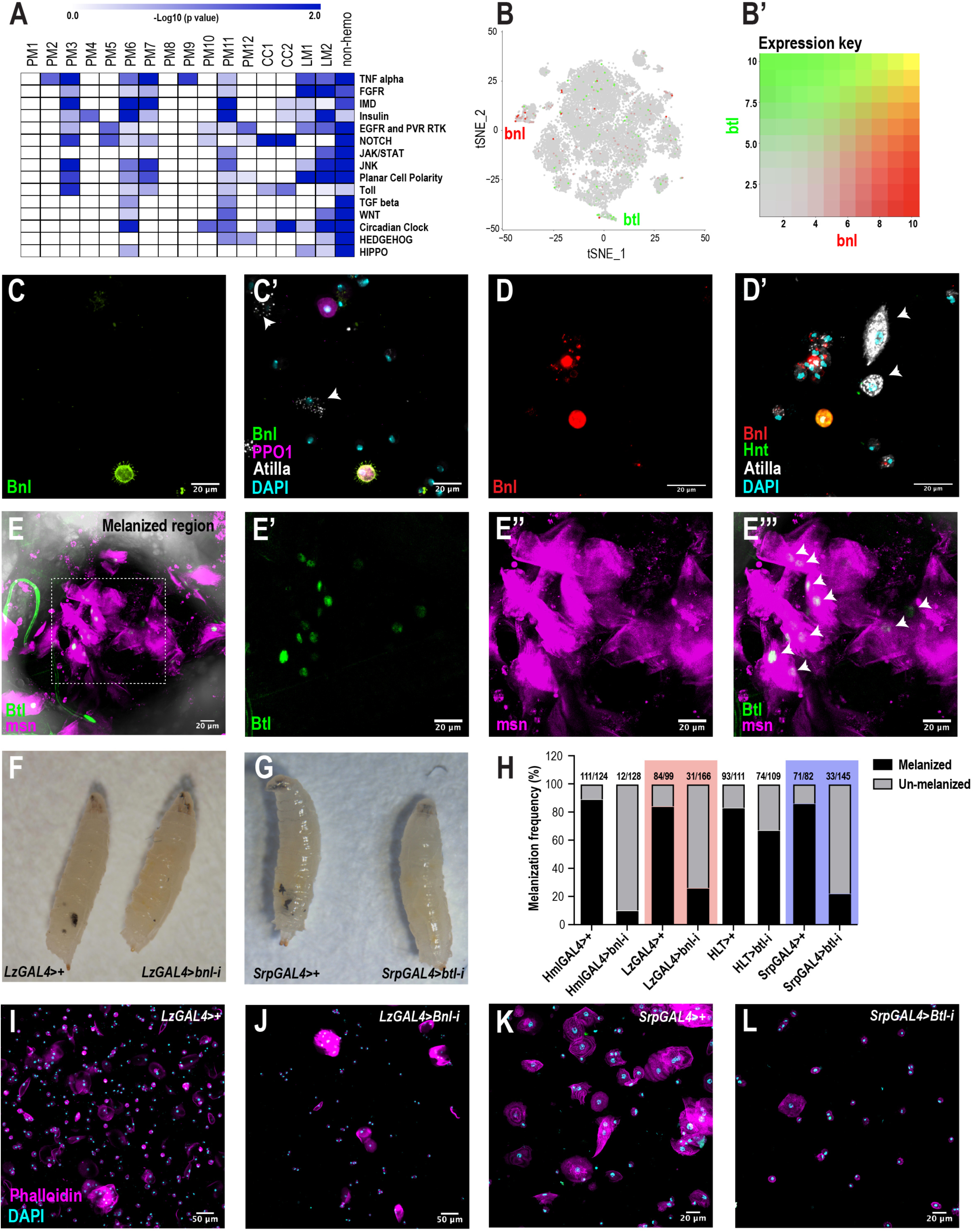
scRNA-seq uncovers a novel role for the FGF pathway in immune response. A) Pathway enrichment of the top marker genes across all the clusters from Fig. 1C. B) Expression of *bnl* (red) and *btl* (green) in CCs and LMs, respectively. B’) Expression heat map key. C, C’) Validation of *bnl* expression in hemocytes of wounded *bnl-lexA; lexAOp-myr-GFP, BcF6-mCherry* larvae. Expression of GFP was detected only in CCs and not LMs (white arrows). GFP and BcF6-mCherry represent the expression of *bnl* and *PPO1*, respectively. Scale bar = 20 μm. D, D’) Validation of *bnl* expression in hemocytes of wasp infested *bnl-lexA; lexAOp-mCherry* larvae. Bnl (mCherry^+^), Bnl+Hnt [Hnt is Hindsight/Peb] (red+green), Atilla (white arrows), DAPI (cyan). Scale bar = 20 μm. E-E’’’) Validation of *btl* expression in LMs *in vivo*. The melanized region of *btlGAL4; UAS-nls-GFP::lacZ, msn-mCherry* larvae was imaged using confocal microscopy. Expression of GFP was detected in LM nuclei (white arrows in E’’’). GFP and msn-mCherry represent the expression of *btl* and *msn*, respectively. *msn* is a marker for LMs. Scale bar = 20 μm. F) Representative images of wasp inf. 48h *LzGAL4>+* (control) and *LzGAL4>bnl-i* larvae. G) Representative images of wasp inf. 48h *SrpGAL4>+* and *SrpGAL4>btl-i* larvae. H) Melanization frequencies of wasp inf. 48h larvae upon *bnl* and *btl* knockdown using *Hml*, *Lz*, *HLT*, and *Srp* GAL4 drivers. I, J) Confocal images of hemocytes from wasp inf. 48h larvae in *LzGAL4>+* controls (I) compared to their *LzGAL4>bnl-i* (J). Scale bar = 50 μm. K, L) Confocal images of hemocytes from wasp inf. 48h larvae in *SrpGAL4>+* controls (K) compared to their *SrpGAL4>btl-i* (L). Scale bar = 20 μm.

To confirm the expression of *bnl* in hemocytes upon wounding, we used *bnl-lexA; lexAOp-GFP, BcF6-mCherry* larvae and found that *bnl* expression was restricted either to CCs or PMs but not LMs (Fig. 6C, C’; S6B). In addition, based on *bnl* expression counts from scRNA-seq, *bnl* may be enriched in PPO1^high^ compared to PPO1^low^ CCs (Fig. S6A). Similar to wounding, we also determined that *bnl* is expressed in subsets of CCs and PMs but not in LMs 48h post infestation (PI) of *bnl-lexA; lexAOp-mCherry* larvae (Fig. 6D, D’). The fraction of Bnl^+^ CCs (as determined by mCherry^+^ Hindsight/Hnt^+^ cells [Hnt/Peb marks CCs]) was very low (∼15.1% ±17.97 standard deviation [SD]; n=20) when compared to the total number of CCs. Next, we examined the expression of *btl* in hemocytes using *btlGAL4; UAS-nls-GFP::lacZ, msn-mCherry*, where *msn-mCherry* marks LMs following wasp infestation (Tokusumi et al., 2009). Confocal imaging at the vicinity of melanized wasp eggs in larvae revealed that a fraction of Btl^+^ LMs were detected near the melanized region, presumably encapsulating the wasp eggs (Fig. 6E-E’’’). Similar to the low fraction of Bnl^+^ CCs, the fraction of Btl^+^ LMs (as determined by GFP^+^ msn^+^ cells) was ∼35% (±12.61 SD, n=5) compared to the total LMs per field of view (Fig. S6E-E’’’). In summary, these results confirm that both *bnl* and *btl* are expressed in rare subsets of CCs and LMs, respectively.

To characterize the role of *bnl* and *btl* in hemocytes, we used the wasp inf. model and knocked down *bnl* in most hemocytes using HmlGAL4. Wasps were allowed to infest second instar *HmlGAL4>control* or *HmlGAL4>bnl-RNAi* (*bnl-i*) larvae. 48h PI, ∼90% of control larvae (111/124) displayed melanotic nodules, which reflect the melanized wasp eggs. In stark contrast, the melanization frequency dropped to <10% (12/128) in *bnl-i* larvae (Fig. 6H), suggesting a defect in melanization. While CC numbers were surprisingly unaltered, the number of all blood cells (DAPI^+^ nuclei), Hml^+^ PMs, and LMs was significantly decreased (Fig. S6F-I). On the other hand, total blood cell number including Hml^+^ PMs, and CCs remained unaltered in uninfested control larvae (Fig. S6F-I). Next, to address the role of *bnl* specifically in CCs, we used LzGAL4 to drive RNAi against *bnl*. Similar to the results obtained with HmlGAL4, over 80% of control larvae (84/99) showed melanized wasp eggs but the melanization frequency dropped to ∼20% (31/166) in *LzGAL4>bnl-i* larvae (Fig. 6F, H), possibly due to the decline in the total blood cell number (Fig. 6I, J; S6J-M). However, the number of CCs remained unchanged and interestingly, while the total number of blood cells and Pxn^+^ hemocytes remained unchanged, CC numbers showed a mild reduction (Fig. S6J-M). These results suggest that *bnl*, derived from CCs, and perhaps a subset of PMs, play a key role in the differentiation and possibly migration of LMs towards wasp eggs for effective melanization.

To address the functions of Btl *in vivo*, we expressed RNAi against *btl* (*btl-i*) using Hml Lineage Tracing (HLT)-GAL4 to maintain GAL4 expression in hemocytes even upon loss of *Hml* expression (*HmlGAL4; UAS-FLP; ubi-FRT-STOP-FRT-GAL4*). After wasp infestation, *HLT-GAL4>btl-i* larvae showed a 67% (74/109) reduction in the melanization frequency compared to that of 83% in controls (93/111) (Fig. 6H). This subtle decline in melanization may be due, in part, to the loss of expression of *Hml* in terminally differentiated LMs and/or possible inefficient flippase activity in Hml^+^ PMs. Thus, we used the pan-hemocyte driver SrpGAL4. *Srp* is well expressed in all blood cells including LMs (Fig. S1G) and as expected, SrpGAL4-mediated knockdown of *btl* revealed a robust decline of ∼20% (31/145) in melanization frequency of 48h post-wasp infested larvae compared to >85% (71/82) in control larvae (Fig. 6G, H). Similar to *bnl-i* in CCs, knockdown of *btl* resulted in a significant reduction in the number of LMs (Fig. 6K, L; S6Q) with a subtle decrease in the total number of blood cells but unchanged CC numbers (Fig. S6N-P) compared to controls. Moreover, none of the cell types, including CCs, displayed any changes in their numbers in uninfested control or *btl-i* larvae (Fig. S6N-P). These data suggest that Btl is critical for the differentiation and possible migration of LMs to elicit an efficient immune response upon wasp infestation. Thus, communication between Bnl^+^ CCs and Btl^+^ LMs may be important in melanization of parasitoid wasp eggs (Fig. S6R).

## Discussion

Previous studies have identified three major *Drosophila* blood cell types essential for combating infections in this species (Banerjee et al., 2019; Lemaitre and Hoffmann, 2007; Rizki, 1957). Here, we used scRNA-seq of larval fly blood to gain deeper insights into the different cell types and their transition states in circulation during normal and inflammatory conditions. Our comprehensive scRNA-seq data provide information on subpopulations of PMs and their immune-activated states. Importantly, our scRNA-seq could precisely distinguish mature CCs and LMs from their respective intermediate states, which are less well understood and for which marker genes were not previously available. Thus, new marker genes identified in this study should facilitate further study of these states. Further, we were able to identify the gene signature of self-renewing PMs and suggest their role as extra-lymph gland oligopotent precursors (Fig. 2J). In addition to the identification of various states of mature cell types, our study also suggests novel roles for a number of genes and pathways in blood cell biology. In particular, we identified a putative new Mtk-like AMP and proposed a role for the FGF signaling pathway in mediating key events leading to the melanization of wasp eggs (Fig. S6R). Finally, we developed a user-friendly searchable online data mining resource that allows users to query, visualize, and compare genes within the diverse hemocyte populations across conditions (Fig. S7A).

### Towards a fly blood cell atlas – defining cell types and states

Blood cell types are dynamic in nature and several transiting intermediate states exist in a continuum during the course of their maturation in several species. Our scRNA-seq analysis provides a framework to distinguish cell types from their various states including oligopotent, transiting intermediate, and activated states.

#### Oligopotent state

Our scRNA-seq analysis identified PM2 as the oligopotent state of PMs based on the enrichment of several cell cycle genes including *polo* and *stg*. This signature suggests that PM2 corresponds to self-renewing PMs located in the circulatory and sessile compartments of the *Drosophila* hematopoietic system where PMs are the only dividing cells identified (Leitão and Sucena, 2015; Makhijani et al., 2011; Rizki, 1957). Further, previous studies suggested that LMs derived from embryonic-lineage hemocytes are readily detectable in circulation prior to their release from the lymph gland (Márkus et al., 2009), and that terminally differentiated CCs can also derive from preexisting PMs in the sessile hub (Leitão and Sucena, 2015). Hence, we propose that PM2 corresponds to the oligopotent state that not only drives expansion of PMs, but importantly can also give rise to CCs and LMs. Our Monocle3 analysis indicates that cell cycle genes decrease over pseudotime and there is ample evidence in support of the notion that cell cycle arrest may be required for terminal differentiation of various cell types in vertebrates (Ruijtenberg and van den Heuvel, 2016; Soufi and Dalton, 2016). Our *in vivo* data also indicates that cell cycle arrest can lead to the generation of terminally differentiated LMs. Interestingly, recent evidence in hemocytes suggests that perturbing cell cycle by knocking down *jumu*, which is upstream of *polo*, can also lead to the generation of LMs by activating Toll (Ahmad et al., 2012; Hao and Jin, 2017; Hao et al., 2018). In contrast, forced expression of certain oncogenes such as activated Ras and Hopscotch/JAK in hemocytes can also lead to overproduction of PMs and LMs (Arefin et al., 2017; Asha et al., 2003; Luo et al., 1995). It is, however, speculated that the proliferation and differentiation of hemocytes in these contexts may be linked to cell cycle (Asha et al., 2003). Thus, it is important to address this paradoxical role of cell cycle in the maintenance of oligopotency and transdifferentiation of PMs. Studies using lineage tracing methods such as G-TRACE (Evans et al., 2009) or CRISPR-based *in vivo* cellular barcoding techniques (Kebschull and Zador, 2018; Spanjaard et al., 2018) may help further characterize the contribution of proliferating oligopotent PMs to blood cell lineages (Fig. 2J).

#### Immune-activated states

PM5 from our scRNA-seq data is enriched in several metabolic genes including *gapdh2* and *lactate dehydrogenase/Ldh*, strongly suggesting that PM5 is metabolically active. It is known that hemocytes can elevate their rates of aerobic glycolysis and increase the levels of *Ldh* to mount an effective immune response against pathogenic bacteria and wasp eggs (Bajgar et al., 2015; Krejčová et al., 2019). Additionally, genes of the glutathione S-transferase family of metabolic enzymes, which are known to catalyze the conjugation of reduced glutathione (GSH) to xenobiotics for their ultimate degradation, are enriched in this state (Fig. S2E). It has been demonstrated that a subset of hemocytes accumulate high GSH levels in *Drosophila* (Tirouvanziam et al., 2004), in support of our data. Further, we classified the two AMP clusters PM6-7 (PM^AMP^) as part of the immune-activated states of PMs. A recent study has demonstrated that AMPs are highly specific and act in synergy against various pathogens (Hanson et al., 2019). Our scRNA-seq analysis reveals the remarkable difference in the expression of a set of AMPs in the two clusters. Future studies with PM^AMP^-specific perturbation of various AMPs identified within PMs should clarify their contribution in killing specific pathogens. Moreover, the role of *Mtkl* against pathogens needs further characterization. Our pseudotime analysis showed that PM^AMP^ ends in the same lineage as LMs suggesting a common mode of activation for these cell types and states. Interestingly, induction in hemocytes of Toll, which is upstream of *Drs*, can lead to the production of LMs (Hao et al., 2018; Schmid et al., 2014), suggesting that LM^int^ cells may act as the common branch point between immune-activated states and LMs.

#### Transiting intermediate states

The transcriptomic composition of CC1 and LM1 clusters suggested the presence of intermediates for CCs and LMs, respectively. Further analysis by Monocle3, which placed these clusters prior to their terminally differentiated cell types, confirmed our hypothesis that CC1 and LM1 correspond to CC^int^ and LM^int^ states, respectively. In the context of the CC lineage within the lymph gland, ultrastructural studies have revealed the presence of immature CCs, called procrystal cells (PCs), alongside mature CCs (Shrestha and Gateff, 1982). We furthered this observation by demonstrating *in vivo* that CCs exist in a continuum (PPO1^low^ to PPO1^high^), validating our Monocle3 and scRNA-seq data. Moreover, clear gene signatures between the CC^int^ and LM^int^ states and their mature counterparts revealed that these intermediates most likely emerge from preexisting Hml^+^ PMs. With regards to the LM lineage, several groups have speculated that intermediates, called podocytes, or also lamelloblasts, may exist based on cell morphology and size (Anderl et al., 2016, 2016; Brantley et al., 2018; Rizki, 1957, 1962). Our scRNA-seq and Monocle3-based data clearly demarcate mature LMs from LM^int^ at the transcriptomic level. In addition, our sub-clustering analysis revealed that LM^int^ possessed a PM signature demonstrating that these intermediates are presumably derived from PM2.

### A novel role for the FGF signaling pathway in hemocyte crosstalk

In addition to the known hemocyte – tissue crosstalk (Shia et al., 2009), *Drosophila* hemocytes must act in a coordinated fashion to combat harmful pathogens and foreign entities such as wasp eggs (Banerjee et al., 2019; Lemaitre and Hoffmann, 2007). However, the signaling pathways that mediate the interactions among hemocytes and wound sites or wasp eggs have been unclear. Our scRNA-seq uncovered a novel role for the FGF signaling pathway in controlling hemocyte differentiation and subsequent effects on the melanization of wasp eggs. The FGF ligand *bnl* and its receptor *btl* were among the genes identified in rare subsets of CCs and LMs, respectively, highlighting the power of scRNA-seq in capturing and detecting these small populations of cells. Based on our *in vivo* data, we propose that Bnl^+^ CCs interact with Btl^+^ LMs to coordinate LM differentiation and possible migration towards parasitoid wasp eggs (Fig. S6R).

In summary, our scRNA-seq data provides a resource for a comprehensive systems-level understanding of *Drosophila* hemocytes across various inflammatory conditions.

## Acknowledgments

We thank Dr. M. Chatterjee, Dr. S. Boswell, and A. Ratner of the single-cell core facility for their kind support in the inDrops-based cell encapsulation. We thank Drs. K. Brückner, S.E. Mohr, J. Zirin, M. Arris, B-E. Campen, R. Vishwanatha, R. Rajakumar, A. Ghosh, P. Saavedra, L. Liu, R. Binari and all members of the Perrimon Lab for their critical comments and helpful insights on the manuscript. We thank Dr. P. Montero Llopis and R. Stephansky of the Microscopy Resources on the North Quad (MicRoN) core facility for their help in imaging, and the *Drosophila* RNAi Screening Center (DRSC) and Bloomington *Drosophila* Stock Center (BDSC) for providing fly RNAi and GAL4 lines used in this study. We also thank Dr. I. Andó for the generous gift of the hemocyte specific antibodies.

NP is an investigator of the Howard Hughes Medical Institute. J.S. is an investigator of the Samsung Science and Technology Foundation under Project Number SSTF-BA1701-15. Work by V.B., M.S., and S.H.S at the Harvard Chan Bioinformatics Core was funded by the Harvard Medical School Tools and Technology Committee and with the support of Harvard Catalyst, The Harvard Clinical and Translational Science Center (NIH award #UL1 RR 025758 and financial contributions from participating institutions). The content is solely the responsibility of the authors and does not necessarily represent the official views of the National Center for Research Resources or the National Institutes of Health. Portions of this research were conducted on the Orchestra High Performance Compute Cluster at Harvard Medical School. This NIH-supported shared facility is partially provided through grant NCRR 1S10RR028832-01.

## Author contributions

S.G.T. and B.C. performed experiments; S.G.T., B.C., Y.H., Y.L., V.B., M.S., S-H. Y., F.L., F.D., R-J.H., J.N., S.H.S., J.S., and N.P. analyzed and discussed the data; A.C. and Y.H. developed the web portal; S.G.T., J.S., and N.P. wrote the manuscript; S.G.T and N.P. conceived and supervised the project.

## Supplemental information

**Figure S1:**
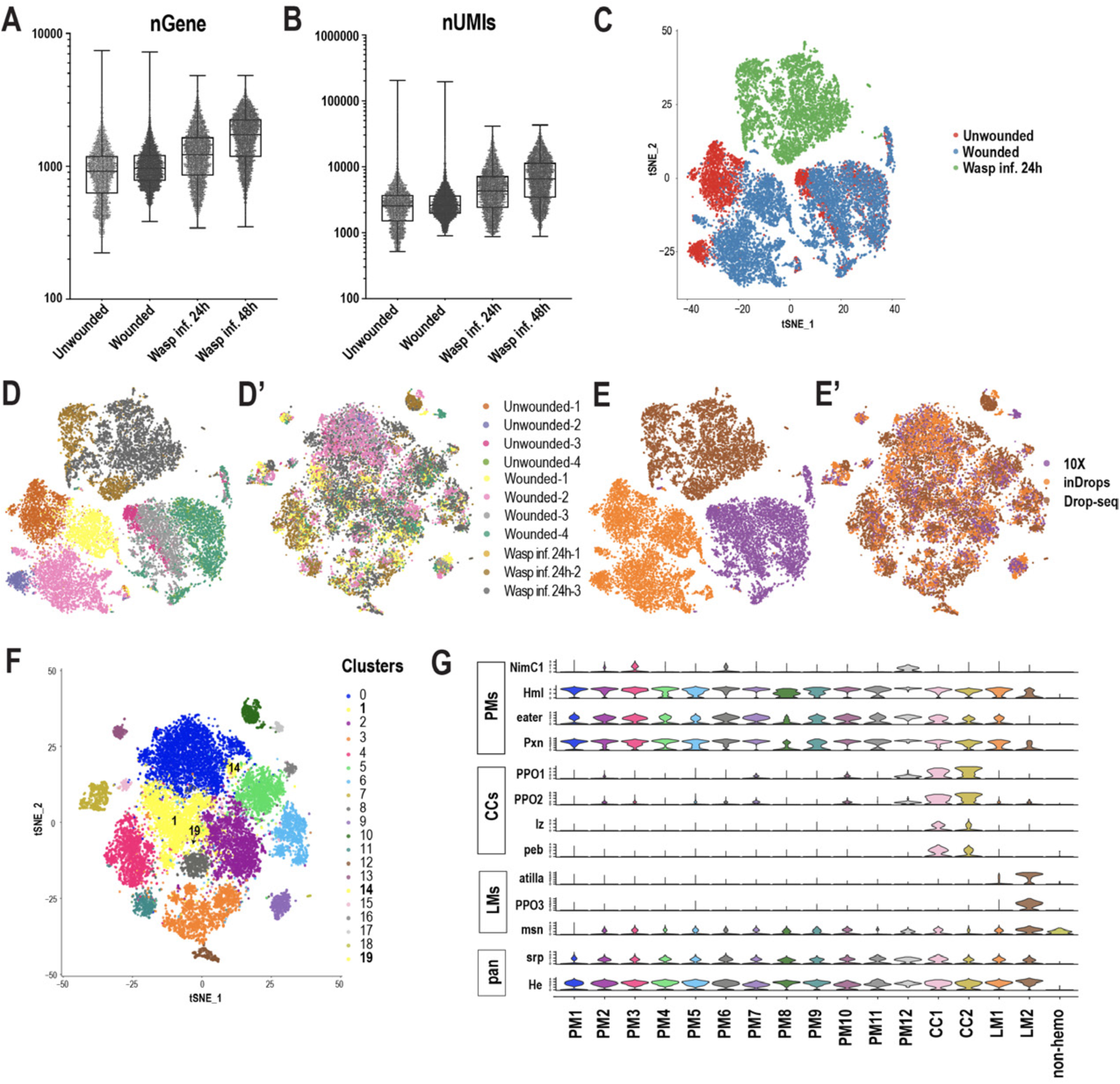
scRNA-seq of *Drosophila* hemocytes. A, B) Number of genes (nGene [A]) and number of unique molecular identifiers (nUMI [B]) detected per cell in Unwounded controls (Unw, n=4), wounded (n=4), wasp infested (wasp inf., 24h, n=3 and wasp inf. 48h, n=3) conditions. C) t-Distributed Stochastic Neighbor Embedding (t-SNE) plot of all the three conditions prior to Harmony-based batch correction and integration of Unw controls (red), Wounded (blue), and Wasp inf. 24h (green) data sets revealed condition specific clusters. Each dot representing one cell is depicted in the t-SNE plot. D, D’) t-SNE plots representing all the replicates across all conditions prior to (D) and after (D’) batch correction. E, E’) t-SNE plots representing all the technologies prior to (E) and after (E’) batch correction. F) t-SNE plot showing all 20 clusters prior to merging clusters 1, 14, and 19 (in yellow) with cluster 0 (see *Merging of clusters* in Methods section for details). G) Violin plots of known marker gene expression to determine the cell types in Fig. 1C

**Figure S2:**
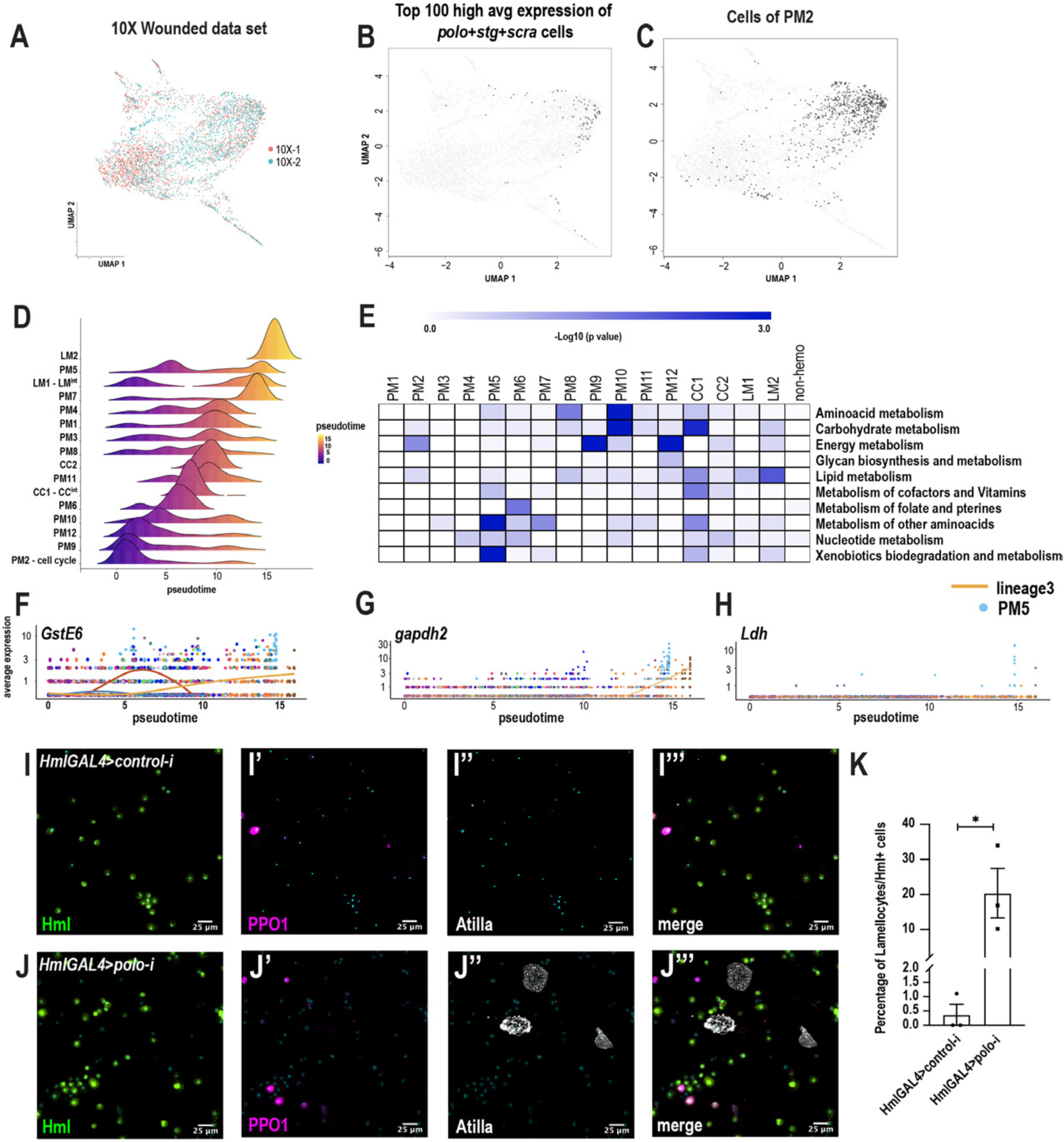
Pseudotemporal ordering of cells delineates immune cell lineages. A) Monocle 3 was used to track cells over pseudotime on the 10X genomics-derived wounded data set. The UMAP represents the two replicates (10X-1 and 10X-2) with ∼4.5K cells. B) UMAP plot represents the cells with high combined average expression of the cell cycle genes *polo*, *stg*, and *scra*. C) UMAP plot shows that the cells with high combined average expression of *polo*, *stg*, and *scra* belong to the PM2 cluster. D) Ridge plots of all the clusters, which suggests that most PMs are PM intermediates. E) Heat map depicts the gene set enrichment pertaining to various metabolic pathways enriched in each cluster (from Fig. 1C). F-H) Gene expression of metabolic genes, such as *GstE6* (F), *gapdh2* (G), and *Ldh* (H), along the pseudotime intervals, reveals that these genes are enriched in PM5 of Lineage 3. I-J’’’) Confocal imaging of hemocytes derived from third instar larvae with HmlGAL4-mediated expression of *control-RNAi* (I-I’’’) or *polo-RNAi* (J-J’’’). Hml^+^ cells, CCs, LMs, and nuclei are represented by EGFP (green), mCherry (PPO1 in magenta), Atilla (white [far red]), and DAPI (cyan), respectively. K) Percentage of LMs normalized to Hml^+^ PMs per field of view in three independent biological replicates (n=3). Error bars are represented as ±SEM (standard error of mean). Statistics were done in Prism using unpaired t-test. P values are represented by * (p<0.05).

**Figure S3:**
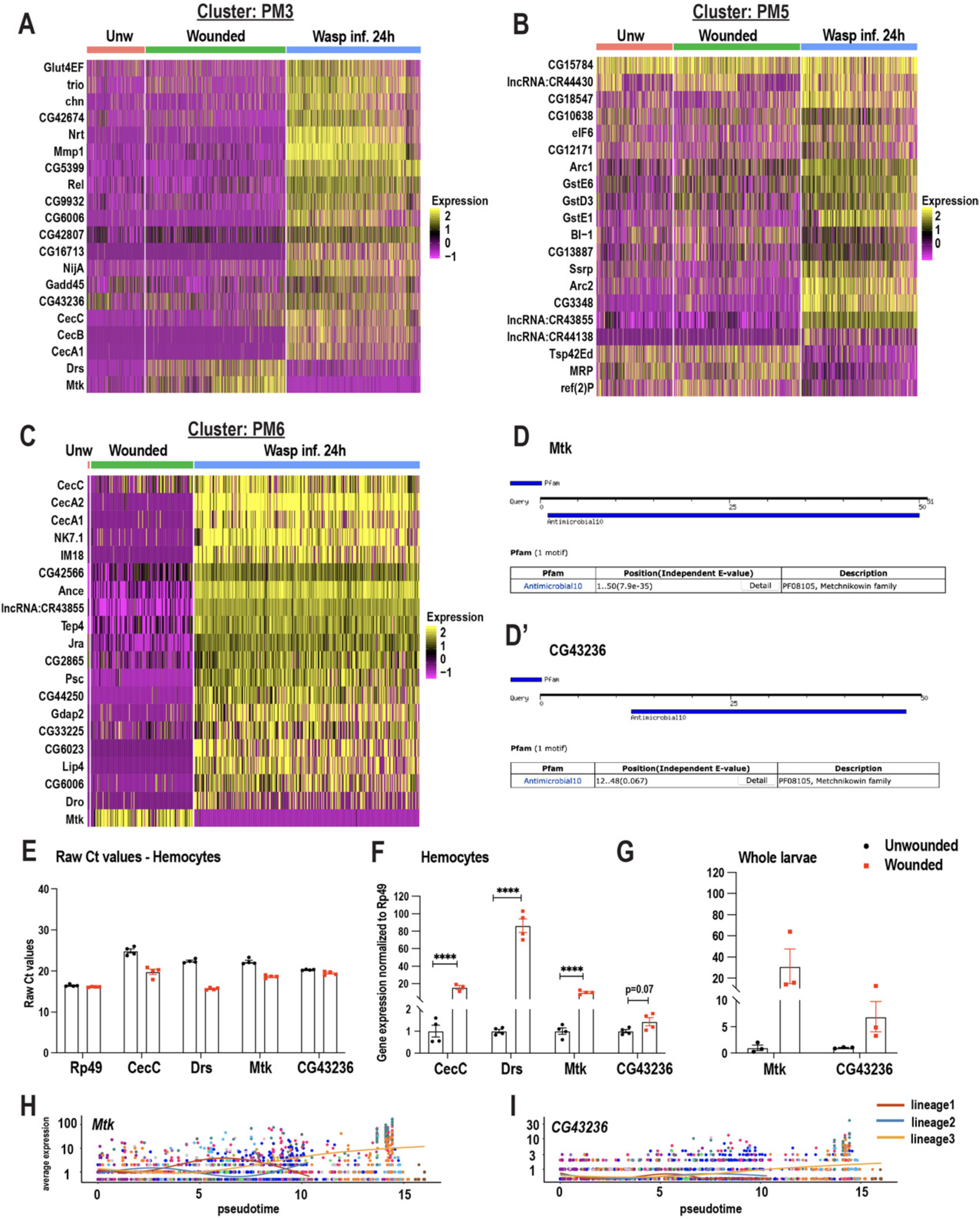
Changes in immune cell composition and identification of a novel Mtk-like AMP. A-C) Heat maps of differentially expressed gene signatures pertaining to PM3 (A), PM5 (B), and PM6 (C). D, D’) Screen shots of the Antimicrobial10 domains within the peptide sequences of Mtk (D) and CG43236 (D’) using the motif search tool https://www.genome.jp/tools/motif/. E) qRT-PCR based raw Ct values corresponding to the AMP genes (*CecC*, *Drs*, *Mtk*, and *CG43236)* demonstrate that *CG43236* is expressed in hemocytes based on the average Ct values [20.3 ± 0.08 (SD) for *CG43236* vs 16.45 ± 0.268 (SD) of *Rp49*] pertaining to Unw control larvae. SD = standard deviation. F) qRT-PCR based relative expression of the AMP genes (*CecC*, *Drs*, *Mtk*, and *CG43236*) in hemocytes derived from unwounded and wounded conditions. G) qRT-PCR based relative expression of the *Mtk* and *CG43236* AMP genes in whole larvae from unwounded or wounded conditions. H-I) Gene expression of *Mtk* (H) and *CG43236* (I) over pseudotime intervals reveals their Lineage3 specific expression. Error bars in E-G are represented as ±SEM (standard error of mean). Statistics were done in Prism using unpaired t-test. P values are represented by * (p<0.05), ** (p<0.01), *** (p<0.001), **** (p<0.0001).

**Figure S4:**
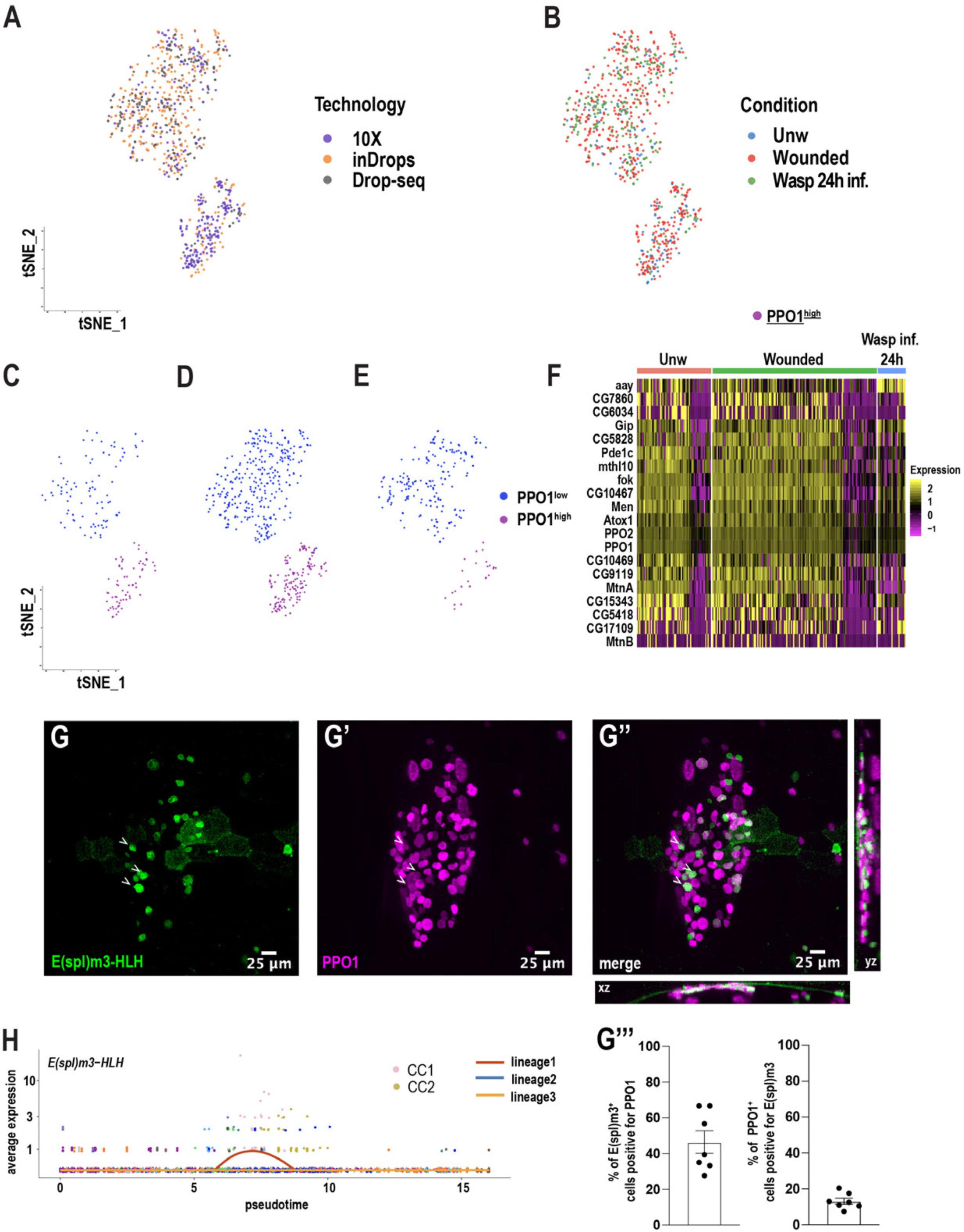
Crystal cell sub-clustering distinguishes CC intermediates from mature CCs. A, B) t-SNE plots represents Harmony-based batch correction per technology (A) and condition (B). C-E) t-SNE plots of unwounded (C), wounded (D), and wasp inf. 24h (E) segregated from the main t-SNE plot pertaining to Fig. 4A. F) Heat map depicting the differentially expressed genes corresponding to the top enriched genes in the PPO1^high^ cluster. Genes were ranked based on expression (logFC) across the different conditions. G-G’’) Expression validation of *E(spl)m3-HLH in vivo.* Confocal images of the CC hub at the posterior-dorsal region of *E(spl)m3-HLH-GAL4; mCD8-GFP, BcF6-mCherry* third instar larvae at steady state reveal that a subset of CCs express *E(spl)m3-HLH*. xz and yz panels in G’’ represent the depth of the confocal Z stacks. G’’’) Percentage of E(spl)m3-HLH^+^ cells positive for PPO1 and percentage of CCs positive for E(spl)m3-HLH. H) Gene expression of *E(spl)m3-HLH* over pseudotime intervals in the CC lineage.

**Figure S5:**
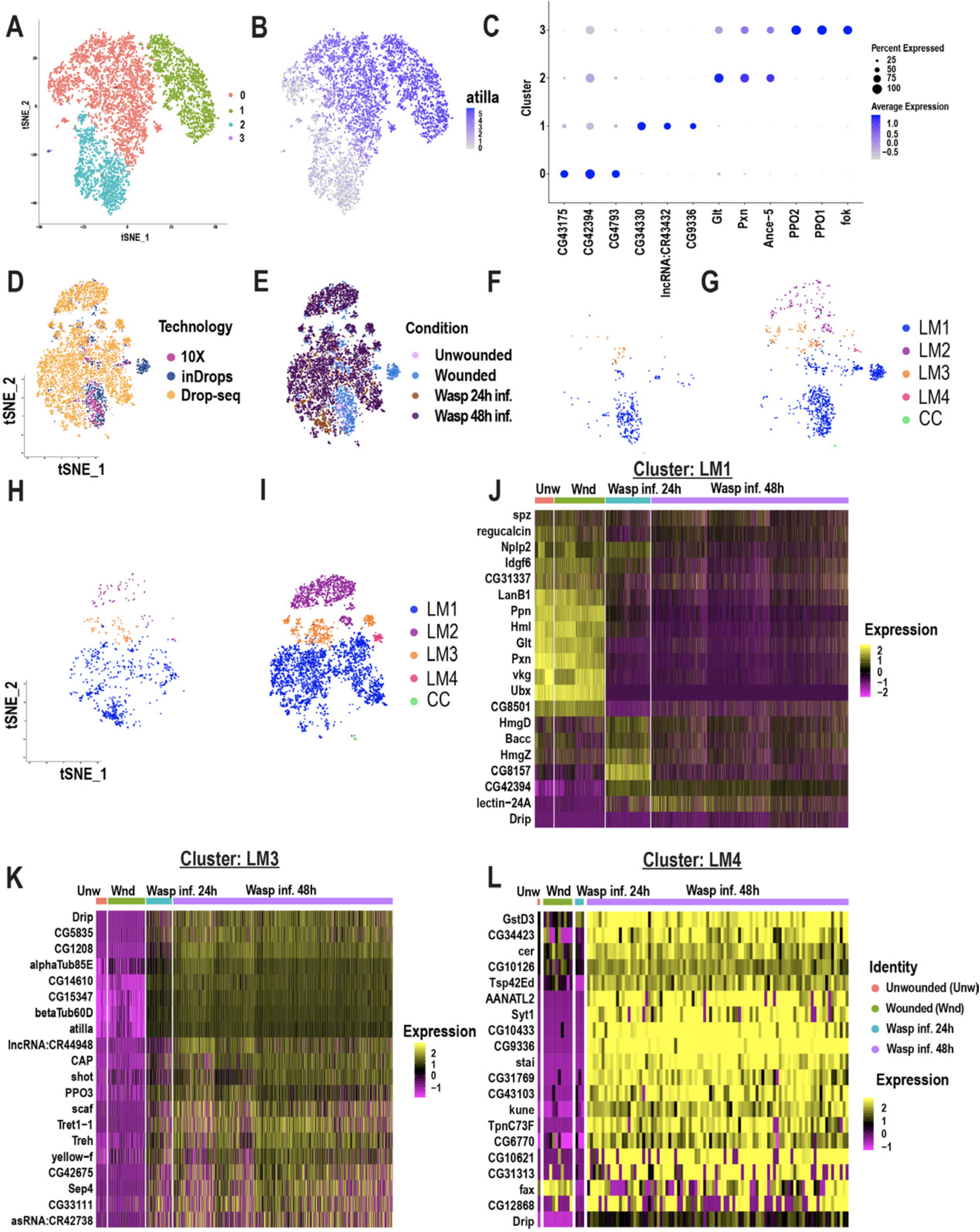
Lamellocyte sub-clustering identifies LM intermediates and subtypes. A) t-SNE plot representing the clustering analysis of wasp inf. 48h data set reveals 4 distinct clusters. B) t-SNE plot demonstrating the expression of *Atilla* in the wasp inf. 48h data set. C) Dot plot shows top enriched genes in each LM sub-cluster. D, E) t-SNE plots represent Harmony-based batch correction per technology (D) and condition (E). F-I) t-SNE plots of unwounded (F), wounded (G), wasp inf. 24h (H), and wasp inf. 48h segregated from the main t-SNE plot pertaining to Fig. 5A. J-L) Heat maps of differentially expressed gene signatures pertaining to the top enriched genes of LM1 (J), LM3 (K), and LM4 (L).

**Figure S6:**
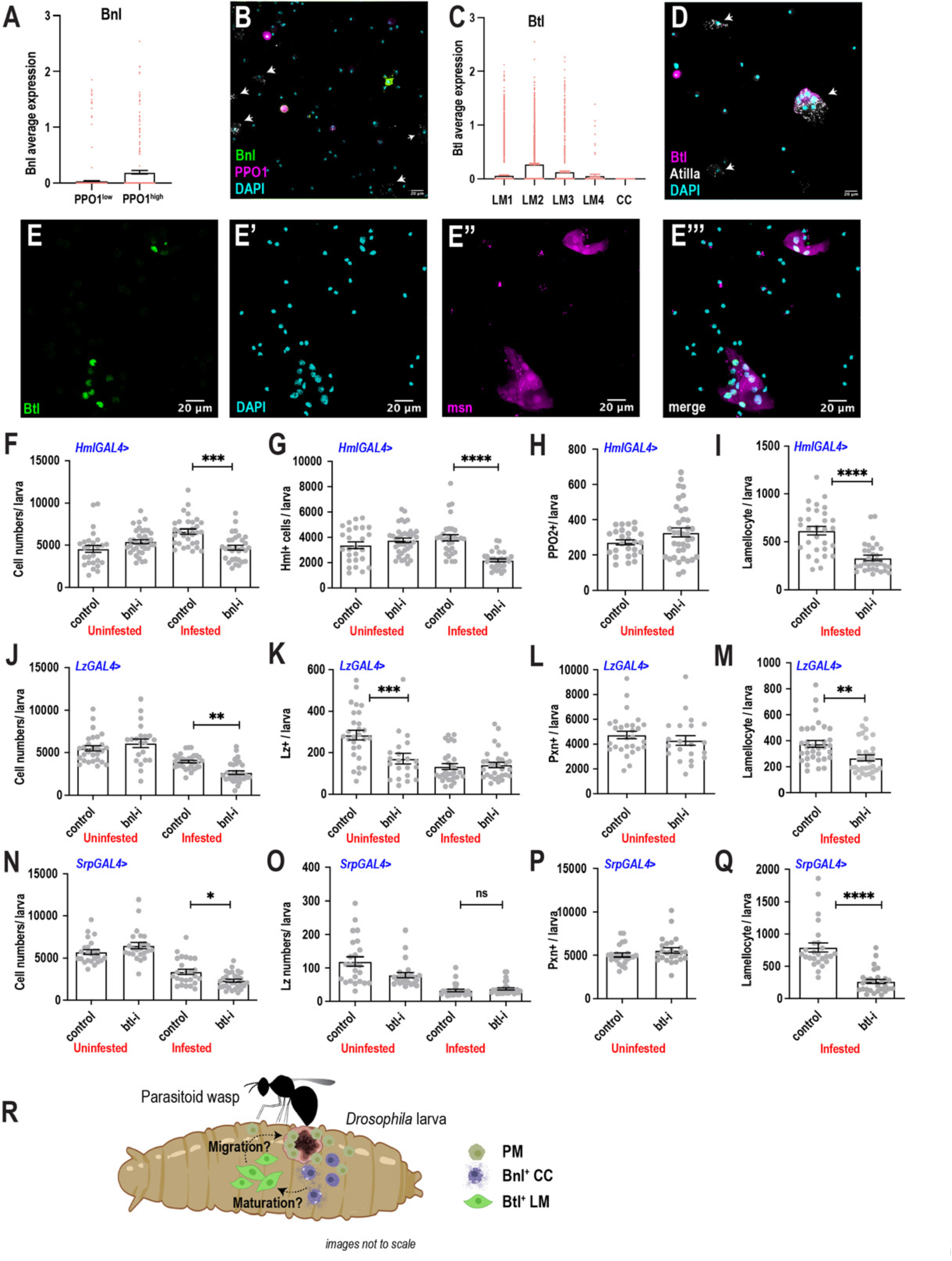
scRNA-seq uncovers a novel role for the FGF pathway in immune response. A) Average *bnl* expression counts derived from the CC sub-clustering data revealed that *bnl* is more enriched in PPO1^high^ CCs. Each colored dot represents one cell. Error bars are represented as ±SEM. B) Expression validation of *bnl* in larval hemocytes upon wounding. Wounded *bnl-lexA; lexAOp-myr-GFP, BcF6-mCherry* larvae were bled and stained for LMs using L1abc antibody. GFP is expressed in CCs and non-CCs but not in LMs (white arrows). C) Average *btl* expression counts derived from the LM sub-clustering data revealed that *btl* is more enriched in LM2 and LM3. Each colored dot represents one cell. Error bars are represented as ±SEM. D) Expression validation of *btl* in larval hemocytes upon wounding. Wounded *btlGAL4; UAS-nls-GFP::lacZ, UAS-mCherry* larvae were bled and stained for LMs using L1abc antibody. GFP is expressed in a subset of LMs (white arrows) and non-LMs. E-E’’’) Validation of *btl* expression in LMs upon wasp infestation. *btlGAL4; UAS-nls-GFP::lacZ, msn-mCherry* larvae were wasp infested and the hemocytes were bled for subsequent staining of the nuclei using DAPI. F-I) Bar graphs representing average number of total cells (F), Hml^+^ cells (G), PPO2^+^ CCs (H), and LMs (I) per larva with or without RNAi against *bnl* using the HmlGAL4 driver. Comparisons were made between uninfested control and wasp inf. 48h conditions. n=26-37 biological replicates per condition and genotype. J-M) Bar graphs representing average number of total cells (J), Lz^+^ CCs (K), Pxn^+^ cells (L), and LMs (M) per larva with or without RNAi against *bnl* using the LzGAL4 driver. Comparisons were made between uninfested control and wasp inf. 48h conditions. n=21-31 biological replicates per condition and genotype. N-Q) Bar graphs representing average number of total cells (N), Lz^+^ CCs (O), Pxn^+^ cells (P), and LMs (Q) per larva with or without RNAi against *btl* using the SrpGAL4 driver. Comparisons were made between uninfested control and wasp inf. 48h conditions. n=24-30 biological replicates per condition and genotype. Error bars in F-Q are represented as ±SEM. Statistics were done in Prism using unpaired t-test. P values are represented by * (p<0.05), ** (p<0.01), *** (p<0.001), **** (p<0.0001). R) Proposed model depicting inter-hemocyte crosstalk between Bnl^+^ CCs and Btl^+^ LMs. Based on our *in vivo* data, we propose that CCs expressing *Bnl* are important for the differentiation or maturation and possible migration of LMs towards parasitoid wasp eggs.

**Figure S7:**
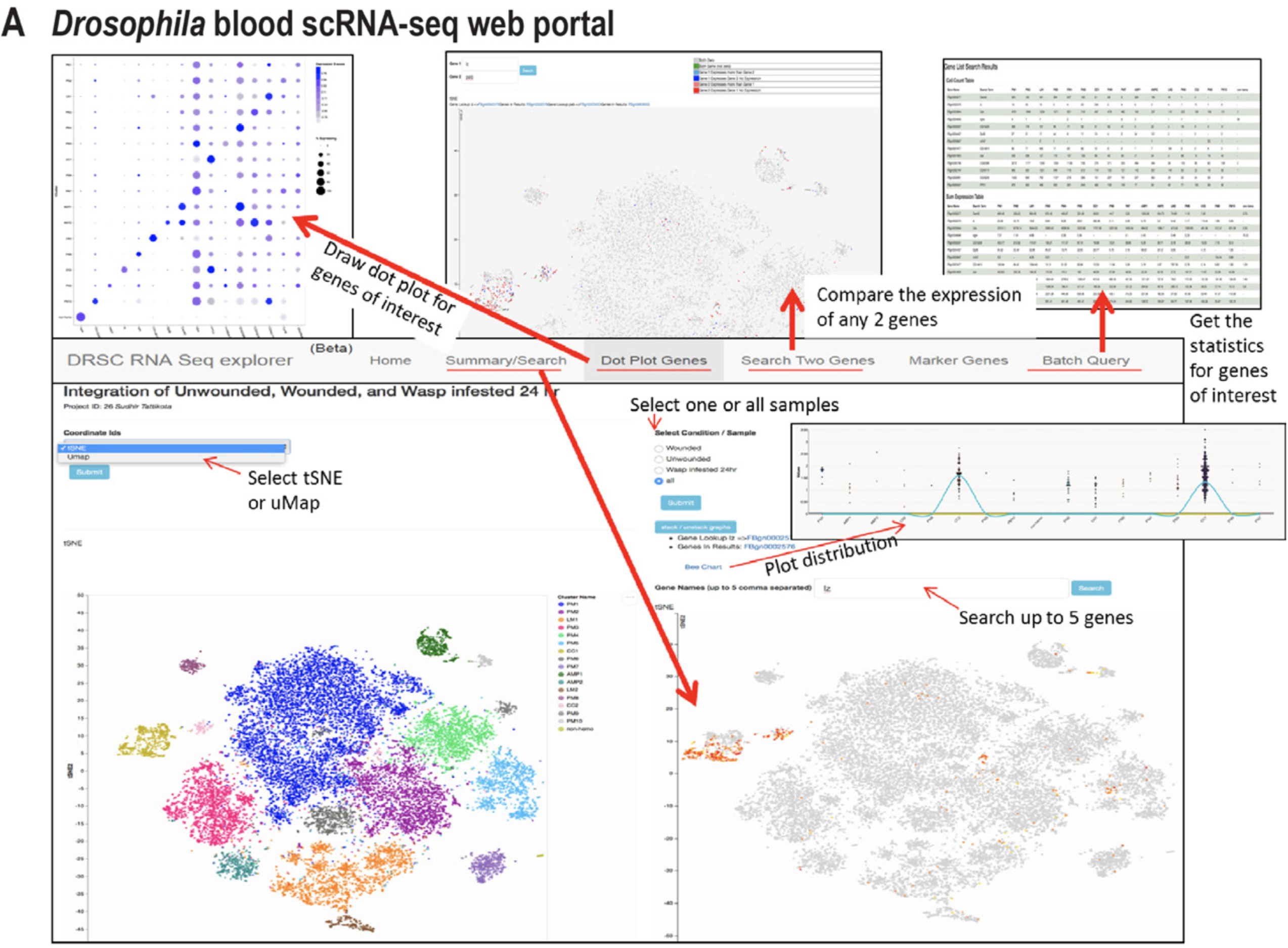
D*r*osophila blood scRNA-seq portal. Snapshot of the searchable *Drosophila* blood scRNA-seq web portal (https://www.flyrnai.org/scRNA/blood/). At the search page, users can search the expression of one gene or the accumulative expression up to 5 genes displayed on the map of choice (t-SNE or UMAP) as well as the sample of choice (unwounded, wounded, wasp infested or all samples together). Users can also view the expression pattern across all the clusters by various plots such as bar graph and violin plot. Users can draw the dot plot for any genes of interest at “Dot Plot Genes” page and compare the expression of any 2 genes at “Search Two Genes” page. The portal allows users to query the markers of any cell cluster of choice as well as getting cluster-based statistics (number of cells’ expression and sum of expression value) at “Batch Query” page.

## Supplementary tables

Table S1: Table representing number of cells, genes, and unique molecular identifiers (UMIs) recovered per sample.

Table S2: Table representing the top marker genes per cluster pertaining to Fig. 1C and D. One cluster per sheet.

Table S3: Table representing the Differentially expressed genes (DEG) per cluster across all conditions pertaining to Fig. 3.

Table S4: Table representing the gene enrichment analysis pertaining to Fig. 6A and S2E.

## Materials and methods

### Fly stocks and reagents

*Drosophila melanogaster* larvae of the genetic backgrounds *w;hmlGAL4Δ, UAS-2X EGFP* (*hml>EGFP*) or *Oregon R* (*OreR*) were used for the preparation of single hemocytes. Third instar *hml>EGFP* larvae and second instar *OreR* larvae were used for wounding and wasp infestations, respectively. To visualize the crystal cell hubs, yw; *lzGAL4; UAS-mCD8::GFP* (*lz>GFP*) (BL# 6314) flies were crossed to *w;;BcF6-mCherry* flies and the resultant female larvae positive for both reporters were used for confocal microscopy. The following stocks were obtained from the Bloomington *Drosophila* Stock Center (BDSC), GAL4 lines: Drip-GAL4 (BL# 66782), E(spl)m3-HLH-GAL4 (BL# 46517), RNAi lines: bnl-RNAi (BL# 34572), Btl-RNAi (BL# 43544), polo-RNAi (BL#33042), luciferase control RNAi (BL# 31603), and Reporter lines: *P{20XUAS-6XmCherry-HA}attP2* (BL# 52268) and *P{13XLexAop2-IVS-myr::GFP}attP40* (BL# 32210). The SrpGAL4 was obtained from a previous study (Waltzer et al., 2003). The *Bnl-LexA* line was a kind gift from Dr. Sougata Roy (Du et al., 2017). The *w;UAS-nlsGFP::lacZ;btlGAL4* line is a Perrimon Lab stock. *BcF6-mCherry* and *msn-mCherry* fly stocks were obtained from Dr. Robert Schulz (Tokusumi et al., 2017).

The *lzGAL4; UAS-mCD8::GFP, BcF6-mCherry* line was obtained by crossing *lzGAL4; UAS-mCD8::GFP* with *BcF6-mCherry* flies. The *Bnl-LexA; LexAOp-myr-GFP, BcF6-mCherry* line (Bnl>GFP; BcF6-mCherry) was obtained by crossing *Bnl-LexA; LexAOp-myr-GFP* and *BcF6-mCherry* flies. The *20X-UAS-6X-mCherry; BtlGAL4, UAS-nlsGFP::lacZ* (Btl>mCherry) line was obtained by crossing *BtlGAL4, UAS-nlsGFP::lacZ* to *20X-UAS-6X-mCherry* reporter line. The *BtlGAL4, UAS-nlsGFP::lacZ, msn-mCherry* line was obtained by crossing *BtlGAL4, UAS-nlsGFP::lacZ* to *msn-mCherry* reporter line.

All flies and larvae were maintained on standard fly food at 25°C.

*Antibodies*: Antibodies against Atilla [anti-mouse L1abc (1:100 dilution)] and PPO1 (1:1000 dilution) were a generous gift from István Ando (Kurucz et al., 2007). Phalloidin (ThermoFischer A34055) was used at a concentration of 1:100. Hindsight (Hnt; 1:10 dilution) antibody was obtained from DSHB.

### Wounding and wasp infestation

#### Wounding

Precisely timed 24h (hours) after egg lay (AEL) larvae of the *hml>EGFP* genotype were collected and grown on normal fly food until they reached 96h AEL for wounding procedure. At 96h AEL, larvae were either left unwounded (Unw) or wounded with a clean tungsten needle (Fine Science Tools, cat# 10130-05). Ten larvae at a time were wounded at their posterior dorsal side and returned back to fly food with a total of at least 80 wounded larvae per vial. 24h later, the Unw control and wounded larvae were retrieved from fly food and washed in distilled water twice.

#### Wasp Infestation

*OreR* larvae were infested at 72h AEL with the wasps of the species *Leptopilina boulardi*. Wasps were removed after 12 hours of co-culture and egg deposition was confirmed by direct observation of wasp eggs in the hemolymph during dissection. 100 larvae were dissected at 96 and 120h AEL, corresponding to 24 and 48h post infestation (wasp inf. 24h and 48h), respectively, in Schneider’s medium (Gibco, cat# 21720024).

### Preparation of single hemocytes in suspension

To get most of the sessile hemocytes into circulation, washed larvae were transferred to 2 ml Eppendorf tubes containing ∼0.5 ml of glass beads (Sigma #9268, size: 425-600 μm) in PBS and larvae were vortexed for 2 min as previously described with minor modifications (Petraki et al., 2015). One set each of Unw and wounded larvae were vortexed in separate tubes at a time. After the brief vortex, larvae were retrieved, washed and transferred to 200 μl of ice-cold PBS in each well of a clean 9-well glass dish per condition. ∼100 larvae were bled by gently nicking open the posterior side of each larva using a pair of clean tweezers. Larvae were allowed to bleed for at least a minute and the hemolymph in PBS was filtered through 100 μm cell strainer and the filtered hemolymph was overlaid onto 2 ml of 1.09 g/ml Optiprep gradient solution (Axis-Shield cat# AXS-1114542) and spun at 2000 rpm for 30 min at 4°C to eliminate dead cells and debris. After centrifugation, ∼150 μl of the hemolymph was transferred to clean low bind Eppendorf tubes and counted using a hemocytometer. High quality single hemocytes were subjected to encapsulation either by inDrops or 10X Genomics v2 platform.

Hemocytes from wasp infested larvae were isolated with some modifications, where the optiprep step was avoided to obtain higher number of cells. Briefly, after vortexing, the larvae were bled in ice cold Schneider’s medium, filtered through 100 μm cell strainer, and transferred to a clean Eppendorf tube. Next, the cells were spun at 4°C for 5 min at 6000 rpm. The supernatant was discarded, and the cells were re-suspended in ice cold PBS to achieve a concentration of 300 cells/μl and subjected to Drop-seq based encapsulation.

### Single hemocyte encapsulation and sequencing

Single hemocytes from Unw control and wounded larvae, respectively, were encapsulated either by inDrops or 10X genomics v2 platforms.

For inDrops, hemocytes were encapsulated at the Single Cell Core facility of the ICCB-Longwood Screening Facility at Harvard Medical School (https://singlecellcore.hms.harvard.edu/) using the inDrops v3 library format (Klein et al., 2015). Reverse transcription and library preparation were performed at the same facility. The libraries were made following a previously described protocol (Klein et al., 2015; Zilionis et al., 2017), with the following modifications in the primer sequences:

RT primers on hydrogel beads: 5’ – CGATTGATCAACGTAATACGACTCACTATAGGGTGTCGGGTGCAG [bc1,8nt] GTCTCGTGGGCTCGGAGATGTGTATAAGAGACAG [bc2,8nt] NNNNNNTTTTTTTTTTTTTTTTTTTV – 3’

R1-N6 primer sequence [step 151 in the library prep protocol (Zilionis et al., 2017)]: 5’ – TCGTCGGCAGCGTCAGATGTGTATAAGAGACAGNNNNNN – 3’

PCR primer sequences (steps 157 and 160 in the library prep protocol in (Zilionis et al., 2017): 5’ – AATGATACGGCGACCACCGAGATCTACACXXXXXXXXTCGTCGGCAGCGTC – 3’, where XXXXXX is an index sequence for multiplexing libraries. 5’ – CAAGCAGAAGACGGCATACGAGATGGGTGTCGGGTGCAG – 3’

With these modifications in the primer sequences, custom sequencing primers are no longer required.

The fragment size of each library was analyzed using a Bioanalyzer high sensitivity chip. Libraries were diluted to 1.5 nM and then quantified by qPCR using primers against the p5-p7 sequence. inDrops libraries were sequenced on an Illumina Nextseq 500 with following parameters: (1) Read 1: 61 cycles, (2) i7 index: 8 cycles index 1, (3) i5 index: 8 cycles (i5), and (4) Read 2: 14 cycles. Binary base call (BCL) files were converted into FASTQ format with bcl2fastq, using the following flags that are required for inDrops v3: (1) --use-bases-mask y*,y*,y*,y*; (2) --mask- short-adapter-reads 0; (3) --minimum-trimmed-read-length 0.

With regards to 10X genomics, cells were encapsulated according to the manufacturer’s protocol. cDNA libraries generated by both platforms were sequenced after pooling four different (indexed) samples per one 400M plus NextSeq500 cartridge with following parameters: (1) Read 1: 26 cycles, (2) i7 index: 8 cycles, (3) i5 index: 0 cycles, and (4) Read 2: 57 cycles.

The protocol for Drop-seq based encapsulation was followed as previously described (Macosko et al., 2015).

### Data processing

#### Count matrix generation

##### inDrops

The software version used to generate counts from the FASTQ files were managed with bcbio-nextgen 1.0.6a0-d2b5b522 (https://github.com/bcbio/bcbio-nextgen) using bioconda (Grüning et al., 2018) (https://bioconda.github.io/). First, cellular barcodes and UMIs were identified for all reads. Second, reads with one or more mismatches of a known barcode were discarded. Third, remaining reads were quasi-aligned to the FlyBase FB2018_02 Dmel Release 6.21 reference transcriptome using RapMap 0.5.0 (Srivastava et al., 2016) (https://github.com/COMBINE-lab/RapMap). EGFP and Gal4 sequences were included in the transcriptome as spike-in genes (https://github.com/hbc/A-single-cell-survey-of-Drosophila-blood). Reads per cell were counted using the umis 0.6.0 package for estimating UMI counts in transcript tag counting data (Svensson et al., 2017) (https://github.com/vals/umis), discarding duplicated UMIs, weighting multi-mapped reads by the number of transcripts they aligned to, and collapsing counts to genes by adding all counts for each transcript of a gene.

Finally, cellular barcodes with fewer than 1,000 reads assigned were discarded from the analysis (see “minimum_barcode_depth” in bcbio documentation for details).

##### 10X genomics

BCL files were analyzed with the Cell Ranger pipeline (v2.1.1). The demultiplexed FASTQ data were aligned to FlyBase FB2018_02 Dmel Release 6.21 reference to generate the single cell count matrix.

##### Drop-seq

Paired-end reads were processed and mapped to the reference genome BDGP 6.02 (Ensembl September 2014) following the Drop-seq Core Computational Protocol version 1.2 (January 2016) and corresponding Drop-seq tools version 1.13 (https://github.com/broadinstitute/Drop-seq) (December 2017) provided by McCarroll Lab (http://mccarrolllab.org/dropseq/). The Picard suite (https://github.com/broadinstitute/picard) was used to generate the unaligned bam files which were processed using the *Drop-seq_alignment.sh* script. The steps include detection of barcode and UMI sequences, filtration and trimming of low-quality bases and adaptors or poly-A tails, and alignment of reads using *STAR* (2.5.3a).

#### Quality control (QC) analysis and filtering

##### inDrops

Gene-level counts were imported to R using the bcbioSingleCell 0.1.15 package (https://github.com/hbc/bcbioSingleCell). This package extends the Bioconductor SingleCellExperiment container class, which is optimized for scRNA-seq (Huber et al., 2015). *SingleCellExperiment: S4 Classes for Single Cell Data*. R package version 1.6.0.] (https://bioconductor.org/packages/SingleCellExperiment). QC analysis was performed using this package, and the ‘filterCells()’ function was used to filter out low quality cells by keeping cellular barcodes with the following metrics: (1) >= 100 UMIs per cell; (2) >= 100 genes per cell; (3) >= 0.85 novelty score, calculated as log10(genes detected) / log10(UMI counts per cell). Additionally, genes with very low expression across the data set were filtered by applying a cutoff of >= 10 cells per gene. One sample, blood3_TCGCATAA (Table S1), was filtered at a higher threshold of 650 genes per cell, which was required to subset the input cellular barcodes into the expected biological range based on the inDrops encapsulation step.

##### 10X genomics

QC was performed by keeping cells with the following metrics: (1) >= 500 UMIs per cell; (2) >= 200 genes per cell; (3) <= 30 percentage of mitochondria genes.

##### DropSeq

Cumulative distribution of reads from the aligned bam files were obtained using *BAMTagHistogram* from the Drop-seq tools package. The number of cells were inferred from the sharp decrease in the slope. The inferred cell number was determined as a minimal threshold number of aligned reads per cell for cell selection.

In our QC pipeline, we did not regress out cell cycle genes during clustering for two reasons: 1. Cell cycle genes did not contribute to the variation for downstream clustering, and 2. We expected cycling PMs to be an important aspect in our analysis. Hence, cell cycle parameters were retained throughout clustering process.

##### Data integration

For combined analysis of all samples, the quality filtered datasets were merged using the common genes into a single Seurat (version 3.1) object (Stuart et al., 2019) and integrated using Harmony (Korsunsky et al., 2019). The gene symbols from inDrops, 10X, and Drop-seq were converted to the same version by using FlyBase online ID Converter tool (FB2019_03) (Thurmond et al., 2019). Genes expressed in at least two out of three conditions were retained when combining datasets from different technologies, to minimize loss of genes, as Harmony uses common genes across all conditions. We ran PCA using the expression matrix of the top 2000 most variable genes. The total number of principal components (PCs) to compute and store were 20. Theta values were set c(2, 10) for condition and technology. A resolution of 0.4 was chosen as clustering parameter. The t-SNE was then performed using default parameters to visualize data in the two-dimensional space.

###### Merging of clusters

At 0.4 resolution, 20 clusters were obtained and three clusters (1, 14, and 19) shared similar gene expression with that of cluster 0. Hence, we merged clusters 1, 14, and 19 into 0 (see Fig. S1F).

###### Sub-clustering

CC sub-clustering and LM sub-clustering follow the same procedure but with different parameters. For CC sub-clustering we used theta=c(2, 5) for condition and technology and clustering resolution of 0.1. For LM sub-clustering we used theta=c(3, 8) for condition and technology and clustering resolution of 0.1.

###### Gene expression visualization by Dot plots

Dot plots were generated using Seurat DotPlot function. Heatmaps were generated using Seurat DoHeatmap function and split by condition.

##### Pseudotemporal ordering of cells using Monocle3

∼4.5K cells from the wounding data set (10X platform; n=2) were analyzed by Monocle3 (https://github.com/cole-trapnell-lab/monocle3) (Cao et al., 2019). The input data set was pre-processed by using num_dim equal 75. Then the data was applied by align_cds function to remove the batch effect. The cell trajectory was calculated by using learn_graph function and ncenter equal 555. Three lineages were selected by using choose_graph_segments function. The gene expression along pseudotime data were extracted from the result of plot_genes_in_pseudotime function. Then the data was used to plot genes along pseudotime in three lineages using ggplot2 v3.2.1 R package and the heatmap was generated using pheatmap v1.0.12 R package. The Ridgeline plot was generated using ggridges v0.5.1 R package.

###### Assignment of the start point

To be unbiased, we calculated the average expression of three cell cycle genes enriched in PM2: *polo*, *stg*, and *scra*. We then assigned the start point, which coincidentally also overlapped with the high expression of *Hml* around the same start point.

#### Data files and analysis code

The original FASTQ files, UMI-disambiguated counts in MatrixMarket Exchange format (MTX files) [see https://math.nist.gov/MatrixMarket/info.html for details], and inDrops v3 sample barcodes used will be available on the Gene Expression Omnibus (GEO submission in progress). The code used to perform clustering and marker analysis is available on GitHub at https://github.com/hbc/A-single-cell-survey-of-Drosophila-blood.

#### Gene set enrichment analysis

We performed gene set enrichment analysis on marker genes with positive fold change for each cluster using a program written in-house. Gene sets for major functional groups were collected from the GLAD database (Hu et al., 2015), and gene sets for metabolic pathways were from the KEGG database (Kanehisa et al., 2017) and Reactome database (McKay and Weiser, 2015). P-value enrichment was calculated based on the hypergeometric distribution. The strength of enrichment was calculated as negative of log10(p-value), which is used to plot the heatmap.

##### Immunostaining and confocal microscopy

###### Whole larval imaging

Third instar *lzGAL4>GFP; BcF6-mCherry* larvae were heat killed in glycerol on a glass slide and directly mounted onto coverslip-bottom imaging dishes (ibidi, cat# 81158). The posterior hemocyte hubs were imaged with Z stacks of 2 μm distance each encompassing the entire area of the hub in a lateral direction using a Zeiss LSM 710 confocal microscope. Finally, all the stacks per larva were merged by summing the intensities on Fiji software for subsequent intensity measurements of GFP and mCherry.

Wasp infested *w;UAS-nlsGFP::lacZ;btlGAL4* larvae were imaged using Nikon C2 Si-plus confocal microscope.

###### Staining hemocytes from unwounded or wounded larvae

∼13 larvae per condition or genotype were vortexed and bled into 300 μl of Schneider’s insect media and then the cells were transferred to one well (per condition) of the chambered coverslip slides. Hemocytes were allowed to settle down for 30 min, then were fixed in 4% paraformaldehyde (PFA) for 20 min. Next, the cells were washed three times with 1X PBS and permeabilized with 0.1% PBST (PBS with 0.1% Triton-X) for 10 min. Subsequently, the cells were blocked with 5% BSA in PBST (blocking buffer) for 20 min and subsequently incubated with primary antibodies L1abc (1:100 dilution) overnight at 4°C. The next day, cells were washed three times in PBST and incubated with corresponding secondary antibodies (1:500 dilution) for 1hr at room temperature. Finally, the cells were washed (3X) and mounting media with DAPI was directly added onto the cells in the wells and imaged using Nikon Spinning disk microscope.

###### Staining hemocytes from wasp infested larvae

Larvae were vortexed with glass beads for one minute before bleeding to detach sessile hemocytes. Next, the larvae were bled on a slide glass (Immuno-Cell Int. cat# 2018A29) and hemocytes allowed to settle onto the slide at 4 °C for 40 minutes. Hemocytes were washed 3 times in 0.4% Triton X-100 in 1x PBS for 10 minutes and blocked in 1% BSA/0.4% TritonX in 1xPBS for 30 minutes. Primary antibody was added, and samples incubated overnight at 4°C. Hemocytes were washed 3 times in 0.4% Triton X in 1xPBS and then incubated with a secondary antibody with 1% BSA/0.4% Triton X in 1xPBS for 3 hours at room temperature. After washing 3 times with 0.4% Triton X in 1xPBS, samples were kept in Vectashield (Vector Laboratory) with DAPI and imaged by a Nikon C2 Si-plus confocal microscope.

###### Cell counting

Hemocytes were mounted and imaged by Nikon C2 Si-plus or Zeiss Axiocam 503. Captured image of hemocytes were quantified and analyzed by ImageJ (plug in: 3D object counter) or Imaris (Bitplane). Hemocytes bled from individual whole larvae were counted for this study.

### Quantitative real time polymerase chain reaction (qRT-PCR)

Hemolymph with hemocytes was derived 24h after wounding along with their Unw control *hml>EGFP* larvae. 50∼80 larvae per biological replicate were bled in 80 μl of Schneider’s insect media and the resulting hemolymph (∼100 μl) was transferred to RNase free Eppendorf tubes and frozen on dry ice. In parallel, whole larvae (4 larvae per biological replicate) were directly transferred to the Eppendorf tubes and frozen on dry ice. For RNA isolation from hemocytes, 1 ml of Trizol was added to each sample and incubated for 5 min before adding 0.2 ml of chloroform for subsequent phase separation. The tubes were spun for 15 min at 12k rpm at 4°C. The aqueous phase was carefully retrieved and transferred to a fresh tube and equal volumes of absolute ethanol was added. For better precipitation of RNA, 1 μl of ultrapure glycogen (1mg/ml) was added to the tubes and incubated at -20°C overnight. Later, the tubes were spun down at 12K rpm for 15 min at 4°C and the resultant pellet was washed in 70% ethanol, air dried, and subjected to DNase treatment using the manufacturer’s instructions of Turbo DNA free (cat# AM1907). The DNA free RNA was further purified using Zymo Direct-zol RNA MicroPrep kit (cat# R2050). The resultant pure RNA was reverse transcribed using BioRad iScript cDNA synthesis kit and SyBr green based qRT-PCR was performed to determine the expression levels of AMP genes. For total RNA isolation from whole larvae, the above protocol was followed except no overnight precipitation method was used and that the larvae were homogenized using RNase-free pestles.

### Statistics

All statistics with regards to intensity measurements, cell counts, and qRT-PCR, were performed on Prism 8 software. The error bars represent ± standard error of mean (SEM) or standard deviation (SD) depending on the sample size as mentioned in the figure legends. Significance between two conditions was calculated by unpaired t test on Prism 8. The p values shown in the figures are represented by * (p<0.05), ** (p<0.01), *** (p<0.001), and **** (p<0.0001).

